# Genomic investigation of the strawberry pathogen *Phytophthora fragariae* indicates pathogenicity is associated with transcriptional variation in three key races

**DOI:** 10.1101/860619

**Authors:** Thomas M. Adams, Andrew D. Armitage, Maria K. Sobczyk, Helen J. Bates, Javier F. Tabima, Brent A. Kronmiller, Brett M. Tyler, Niklaus J. Grünwald, Jim M. Dunwell, Charlotte F. Nellist, Richard J. Harrison

## Abstract

The oomycete *Phytophthora fragariae* is a highly destructive pathogen of cultivated strawberry (*Fragaria* × *ananassa*), causing the root rotting disease, ‘red core’. The host-pathogen interaction has a well described gene-for-gene resistance relationship, but to date neither candidate avirulence nor resistance genes have been identified. We sequenced a set of American, Canadian and UK isolates of known race type, along with three representatives of the closely related pathogen of the raspberry (*Rubus idaeus*), *Phytophthora rubi*, and found a clear population structure, with a high degree of nucleotide divergence seen between some race types and abundant private variation associated with race types 4 and 5. In contrast, between isolates defined as UK races 1, 2 & 3 (UK1-2-3) there was no evidence of gene loss or gain; or the presence of insertions/deletions (INDELs) or Single Nucleotide Polymorphisms (SNPs) within or in proximity to putative pathogenicity genes could be found associated with race variation. Transcriptomic analysis of representative UK1-2-3 isolates revealed abundant expression variation in key effector family genes associated with pathogen race; however, further long read sequencing did not reveal any long range polymorphisms to be associated with avirulence to race UK2 or UK3 resistance, suggesting either control in *trans* or other stable forms of epigenetic modification modulating gene expression. This work reveals the combined power of population resequencing to uncover race structure in pathosystems and *in planta* transcriptomic analysis to identify candidate avirulence genes. This work has implications for the identification of putative avirulence genes in the absence of associated expression data and points towards the need for detailed molecular characterisation of mechanisms of effector regulation and silencing in oomycete plant pathogens.

## INTRODUCTION

*Phytophthora fragariae*, the causal agent of red core or red stele root rot, is a highly destructive pathogen of cultivated strawberry (*Fragaria* × *ananassa*), resulting in whole plant collapse. The majority of commercial strawberries grown in the UK are grown on table tops using soilless substrate, under polytunnels or in glasshouses (Robinson Boyer et al., 2016). *Phytophthora* spp. are a particular problem in these systems due to the ease of spread through the irrigation system via the motile zoospores. Since the first report in Scotland in 1920, this disease has spread to the majority of strawberry growing regions, except China and the Southern Mediterranean regions of Europe (van de Weg, 1997b; EFSA Panel on Plant Health (PLH), 2014). Currently, it is treated as a quarantine pest by the European and Mediterranean Plant Protection Organization (EPPO), where it is listed as an A2 pest (van de Weg, 1997b; EPPO, 2018). The classification of this pathogen has proven controversial, as initially the organism was identified as a single species (Hickman, 1941), but when a *Phytophthora* disease of raspberry (*Rubus idaeus*) was discovered, it was reclassified as *P. fragariae* var. *fragariae* (Wilcox et al., 1993). More recently, the pathogens have been separated into distinct species, *P. fragariae* and *Phytophthora rubi*, affecting strawberry and raspberry respectively. This was supported by sequence analysis of key loci (Man in ’t Veld, 2007), as well as whole genome analyses (Tabima et al., 2018).

It has previously been proposed that the ability of different isolates of *P. fragariae* to cause disease on a variety of *F.* × *ananassa* cultivars can be explained by a gene-for-gene model (van de Weg, 1997a). The model is currently thought to consist of at least eleven resistance genes in *F.* × *ananassa* with eleven corresponding avirulence factors in *P. fragariae* (W. E. van de Weg, Wageningen University and Research, The Netherlands, personal communication). The development of race schemes is country dependent and ones exist for the UK, USA and Canada.

All publicly available genome assemblies of *P. fragariae* have to date solely utilised Illumina short read sequencing technologies, resulting in assemblies of 73.68 Mb and 76 Mb, in 1,616 and 8,511 scaffolds respectively (Gao et al., 2015; Tabima et al., 2017). Recently, long read sequencing technology has been shown to provide assemblies of improved contiguity for *Phytophthora* pathogens, specifically the generation of the haplotype-phased assembly of *Phytophthora ramorum* (60.5 Mb in 302 primary contigs) using PacBio sequencing (Malar et al., 2019) and the assembly of *Phytophthora capsici* (95.2 Mb in 424 scaffolds) using Oxford Nanopore Technology (Cui et al., 2019).

Pathogenomic investigations in *Phytophthora* species of pathosystems with similar gene-for-gene models of resistance have shown a variety of mechanisms through which variation in virulence can be controlled. For instance, in *Phytophthora sojae*, the *Avr1d* gene was identified as an RxLR effector recognised by the *Rps1d* resistance gene in soybean (Yin et al., 2013). Studies of the RxLR effector *PiAvr4* from *Phytophthora infestans* showed that it was always present in isolates avirulent on potato plants containing the resistance gene *R4*, whereas virulent isolates possessed a frameshift mutation producing a truncated protein (van Poppel et al., 2008). In comparison, *Avr3c* in *P. sojae* was identified in both virulent and avirulent isolates on soybean plants containing *Rps3c*, but in virulent isolates the gene displayed several polymorphisms resulting in a change to the amino acid sequence leading to a failure of recognition by the plant (Dong et al., 2009). Recently, investigations of the EC-1 clonal lineage of *P. infestans* revealed a variation of the ability of isolates to cause disease on potato plants possessing the *Rpi-vnt1.1* gene. It was shown, in the absence of genetic mutations, that differences in the expression level of *Avrvnt1* were detected and these correlated with virulence (Pais et al., 2018).

In this study, we assembled and annotated a population of isolates of *P. fragariae* and the related pathogen of raspberry, *P. rubi*. We identified a subpopulation of *P. fragariae* isolates representing three distinct pathogenicity races (UK1-2-3). This subpopulation was found to be remarkably similar in gene complement, as well as showing little divergence at the nucleotide sequence level. To further investigate the cause of the observed variation in pathogenicity phenotypes, transcriptomic datasets were generated for representative isolates of each pathogenicity race in this subpopulation. This revealed expression level polymorphisms between the isolates, allowing for the generation of candidate lists for *PfAvr1*, *PfAvr2* and *PfAvr3*. A strong candidate for *PfAvr2*, PF003_g27513 was identified as expressed in the BC-16 and A4 (UK2) isolates, yet not expressed in the BC-1 (UK1) and NOV-9 (UK3) isolates. A candidate for *PfAvr3* was also identified, PF009_g26276; it is expressed in NOV-9 (UK3), yet not expressed in BC-1 (UK1), A4 (UK2) and BC-16 (UK2) isolates. Additional sequencing did not reveal long-range polymorphisms influencing the expression of these candidate genes; we therefore suggest control may be in *trans* or due to epigenetic factors.

## MATERIALS AND METHODS

### Isolate selection and sources

A selection of ten isolates of *P. fragariae* were sourced from the Atlantic Food and Horticulture Research Centre (AFHRC), Nova Scotia, Canada. An additional isolate of *P. fragariae*, SCRP245, alongside three *P. rubi* isolates, were sourced from the James Hutton Institute (JHI), Dundee, Scotland (detailed in **Table 1**).

**TABLE 1.** | **Summary of Phytophthora fragariae and Phytophthora rubi isolates used in this study.**

| Isolate Name | Species | Location | Date Isolated | Isolated by | Pathogenicity Race |
| --- | --- | --- | --- | --- | --- |
| A4 | *Phytophthora fragariae* | Unknown | 17/12/2001 | Unknown | US4 |
| BC-1 | *Phytophthora fragariae* | Commercial Strawberry Field, Delta, BC, Canada | 05/01/2007 | N. L. Nickerson | CA1 (UK1) |
| BC-16 | *Phytophthora fragariae* | Commercial Strawberry Field, Ladner, BC, Canada | 05/01/2007 | N. L. Nickerson | CA3 (UK2) |
| BC-23 | *Phytophthora fragariae* | Commercial Strawberry Field, Aldergrove, BC, Canada | 30/01/2012 | N. L. Nickerson | CA5 |
| NOV-5 | *Phytophthora fragariae* | Commercial Strawberry Field, Nine Mile River, Hants County, NS, Canada | 17/12/2001 | N. L. Nickerson | CA1 |
| NOV-9 | *Phytophthora fragariae* | Commercial Strawberry Field, Billtown, Kings County, NS, Canada | 05/01/2007 | N. L. Nickerson | CA2 (UK3) |
| NOV-27 | *Phytophthora fragariae* | Commercial Strawberry Field, Cambridge Station, Kings County, NS, Canada | 19/12/2001 | N. L. Nickerson | CA2 |
| NOV-71 | *Phytophthora fragariae* | Commercial Strawberry Field, Middle Clyde River, Shelburne County, NS, Canada | 05/01/2007 | N. L. Nickerson | CA2 |
| NOV-77 | *Phytophthora fragariae* | Commercial Strawberry Field, Nine Mile River, Hants County, NS, Canada | 30/01/2012 | N. L. Nickerson | CA5 |
| ONT-3 | *Phytophthora fragariae* | Commercial Strawberry Field, Fort Erie, ON, Canada | 05/01/2007 | N. L. Nickerson | CA4 |
| SCR245 | *Phytophthora fragariae* | Kent, England, UK | 1945 | Unknown | Unknown |
| SCR249 | *Phytophthora rubi* | Germany | 1985 | Unknown | 1 |
| SCR324 | *Phytophthora rubi* | Scotland, UK | 1991 | Unknown | 1 |
| SCR333 | *Phytophthora rubi* | Scotland, UK | 1985 | Unknown | 3 |

### Culturing of isolates

All work with *P. fragariae* and *P. rubi* was conducted in a Tri-MAT Class-II microbiological safety cabinet. Isolates were routinely subcultured on kidney bean agar (KBA) produced as previously described (Maas, 1972). Isolates were grown by transferring two pieces of between 1 and 4 mm^2^ onto fresh KBA plates, subsequently sealed with Parafilm^®^. Plates were then grown at 20°C in the dark in a Panasonic MIR-254 cooled incubator for between seven and fourteen days.

Mycelia were also grown in liquid pea broth, produced as previously described (Campbell et al., 1989) with the addition of 10 g/L sucrose. These plates were inoculated with five pieces (1 – 4 mm^2^) of mycelium and media, subsequently sealed with Parafilm^®^. These were grown at 20 °C for four to five days in constant darkness.

### Pathogenicity testing of isolates

Mother stock plants of *F*. × *ananassa* were maintained in 1 L pots in polytunnels. Runner plants were pinned down into 9 cm diameter pots filled with autoclaved 1:1 peat-based compost:sand. The clones were grown on for three weeks to establish their own root system and then were cut from the mother plant. Inoculations were performed as described previously (van de Weg et al., 1996). Plants were placed in a growth chamber with 16/8 hour light/dark cycle at a constant 15 °C. Inoculated plants stood in a shallow layer of tap water (2 – 7 mm) for the entire experiment and were watered from above twice a week. After six weeks, plants were dug up and the roots were rinsed to remove the soil/sand mix. Roots were then assessed for distinctive disease symptoms of ‘rat’s tails’, which is the dieback from the root tip and ‘red core’, which is the red discolouration of the internal root visible when longitudinally sliced open. Samples for which infection was unclear were visualised under a light microscope for the presence of oospores. This was performed through squash-mounting of the root tissue, where a sample of root was excised and pressed between a microscope slide and cover slip. This sample was then examined at 40x magnification under high light intensity with a Leitz Dialux 20 light microscope.

### Sequencing of DNA

For Illumina sequencing, gDNA was extracted from 300 mg of freeze-dried mycelium using the Macherey-Nagel NucleoSpin^®^ Plant II Kit. The manufacturer’s protocol was modified by doubling the amount of lysis buffer PL1 used, increasing the incubation following RNase A addition by five minutes, doubling the volume of buffer PC and eluting in two steps with 35 µL of buffer PE warmed to 70 °C. For PacBio and Oxford Nanopore Technologies (ONT) sequencing, gDNA was extracted using the Genomic-Tip DNA 100/G extraction kit, following the Tissue Sample method.

To create Illumina PCR-free libraries, DNA was sonicated using a Covaris M220 and size-selected using a BluePippin BDF1510 1.5% gel selecting for 550 bp. Libraries were constructed using the NEBNext enzymes: End repair module (E6050), A-TAiling module (E6053), Blunt T/A ligase (M0367) and Illumina single-indexed adapters. Sequencing was performed to generate 2 × 300 bp reads on a MiSeq^TM^ system using MiSeq Reagent Kit V3 600 cycle (MS-102-3003). PacBio library preparation and sequencing was performed by the Earlham Institute, UK on a PacBio RS II machine. ONT sequencing libraries were created using the SQK-LSK108 kit following the manufacturer’s protocol and sequenced using the FAH69834 FLO-MIN106 flow cell on a GridION for approximately 28 hours.

### Inoculation time course experiment

Growth of isolates for inoculations were performed on fresh KBA plates for approximately 14 days as described above, before the mycelium had reached the plate edge. Plugs of mycelium growing on agar were taken using a flame sterilised 10 mm diameter cork borer and plugs were submerged in chilled stream water. Plates were incubated in constant light for three days at 13 °C, with the water changed every 24 hours. For the final 24 hours, chilled Petri’s solution (Judelson et al., 1993) was used. Roots of micropropagated *F.* × *ananassa* ‘Hapil’ plants (GenTech, Dundee, UK), maintained on *Arabidopsis thaliana* salts (ATS) media (Taylor et al., 2016), were submerged for one hour before transfering back to ATS plates. These plates were kept at 15 °C, with 16/8 hour light/dark cycle, with a photosynthetic photon flux (PPF) of 150 µmol m^−2^ s^−1^ provided by fluorescent lamps (FL40SSENW37), in a Panasonic MLR-325H controlled environment chamber. Root tissue was harvested at a selection of time points post inoculation by rinsing root tissue successively in three beakers of sterile dH2O to remove all media. Roots were separated below the crown tissue, flash frozen in liquid nitrogen and stored at −80 °C.

Extraction of total RNA from inoculated root tissue was performed similarly to the previously described 3% CTAB3 method (Yu et al., 2012). Briefly, plant material was disrupted under liquid nitrogen in a mortar and pestle, previously decontaminated through cleaning with RNaseZap^TM^ solution and baking for 2 hours at 230 °C to deactivate RNAse enzymes. This material was transferred to a warmed extraction buffer (Yu et al., 2012) containing β-mercaptoethanol with 0.01 g of PVPP added per 0.1 g of frozen tissue. The remaining steps were performed as previously described (Yu et al., 2012), except that 60 µL DEPC-treated H2O was used to elute RNA.

Mycelium was grown in liquid pea both and dried on a Q100 90 mm filter paper (Fisher Scientific) using a Büchner funnel and a Büchner flask attached to a vacuum pump. Total RNA was then extracted using the QIAGEN RNEasy Plant Mini kit with the RLC buffer. Extraction was carried out following the manufacturer’s instructions with an additional spin to remove residual ethanol.

RNA was checked for quality using a NanoDrop 1000 spectrophotometer, quantity with a Qubit 2.0 Fluorometer and integrity with a TapeStation 4200 before being prepared for sequencing via Reverse Transcription-Polymerase Chain Reaction (RT-PCR) and sequencing on an Illumina HiSeq^TM^ 4000 by Novogene, Hong Kong, Special Administrative Region, China. Timepoints for sequencing were selected through the detection of β-tubulin transcripts by RT-PCR. Reverse transcription was performed with the SuperScript^TM^ III Reverse Transcriptase kit with an equal amount of RNA used for each sample. The complementary DNA (cDNA) was then analysed by PCR with 200 µM dNTPs, 0.2 µM of each primer (detailed in Supplementary Table S1), 2 µL of cDNA template and 2.5 units of Taq DNA polymerase and the buffer supplied in a 20 µL reaction. Reactions were conducted in a Veriti 96-well thermocycler with an initial denaturation step at 95 °C for 30 seconds, followed by 35 cycles of a denaturation step at 95 °C for 30 seconds, an annealing temperature of 60 °C for 30 seconds and an extension step of 72 °C for 30 seconds. This was followed by a final extension step of 72 °C for 5 minutes and held at 10 °C. Products were visualised by gel electrophoresis on a 1% w/v agarose gel at 80 V for 90 minutes, stained with GelRed. Following this, three biological replicates of samples taken at: 24 hpi, 48 hpi and 96 hpi for BC-16, 48 hpi for BC-1 and 72 hpi for NOV-9 were sequenced (Supplementary Figure S1). Additionally, *in vitro* mycelial RNA was sequenced.

*P. rubi* RNA-Seq reads were sequenced on a HiSeq^TM^ 2000 system.

### Genome assembly

Prior to assembly, Illumina reads were cleaned and sequencing adaptors were removed with fastq-mcf (Aronesty, 2013). Quality control of PacBio data was performed by the Earlham Institute, Norwich, UK. ONT reads were basecalled with Albacore version 2.2.7 and adaptors were removed with Porechop version 0.2.0 (Wick, 2018) and the trimmed reads were corrected with Canu version 1.4 (Koren et al., 2017).

Assemblies of isolates sequenced solely with Illumina data were generated with SPAdes version 3.11.0 (Bankevich et al., 2012) with Kmer sizes of 21, 33, 55, 77, 99 and 127. PacBio reads were assembled using FALCON-Unzip (Chin et al., 2016). FALCON version 0.7+git.7a6ac0d8e8492c64733a997d72a9359e1275bb57 was used, followed by FALCON-Unzip version 0.4.0 and Quiver (Chin et al., 2013). Error corrected ONT reads were assembled using SMARTdenovo version 1.0.0 (Ruan, 2018).

Following assembly, error correction was performed on ONT assemblies by aligning the reads to the assembly with Minimap2 version 2.8r711-dirty (Li, 2018) to inform ten iterations of Racon version 1.3.1 (Vaser et al., 2017). Following this, reads were again mapped to the assembly and errors were corrected with Nanopolish version 0.9.0 (Simpson, 2018). PacBio and ONT assemblies had Illumina reads mapped with Bowtie 2 version 2.2.6 (Langmead and Salzberg, 2012) and SAMtools version 1.5 (Li et al., 2009) to allow for error correction with ten iterations of Pilon version 1.17 (Walker et al., 2014).

Following assembly, all contigs smaller than 500 bp were discarded and assembly statistics were collected using Quast version 3.0 (Gurevich et al., 2013). BUSCO statistics were collected with BUSCO version 3.0.1 (Simão et al., 2015) using the eukaryota_odb9 database on the assemblies, as the stramenopile database was not available at the time of the analysis. Identification of repetitive sequences was performed with Repeatmasker version open-4.0.5 (Smit et al., 2015) and RepeatModeler version 1.73 (Smit and Hubley, 2015). Transposon related sequences were identified with TransposonPSI release 22^nd^ August 2010 (Haas, 2010).

### Prediction of gene models and effectors

Gene and effector prediction was performed similarly to a previously described method (Armitage et al., 2018). Firstly, raw RNA-Seq reads of BC-16 from both mycelial samples and the inoculation time course were cleaned with fastq-mcf (Aronesty, 2013). Reads from the inoculation time course were first aligned to the *Fragaria vesca* version 1.1 genome (Shulaev et al., 2011) with STAR version 2.5.3a (Dobin et al., 2013) and unmapped reads were kept. These unmapped reads and those from *in vitro* mycelium were mapped to the assembled *P. fragariae* genomes with STAR (Dobin et al., 2013). RNA-Seq data from *P. rubi* were aligned to *P. rubi* assemblies with the same method. Further steps were performed as previously described (Armitage et al., 2018). Additionally, putative apoplastic effectors were identified through the use of ApoplastP (Sperschneider et al., 2018). Statistical significance of the differences between predicted numbers of effector genes was assessed with a Welch two sample t-test in R version 3.4.3 (R core team, 2017).

### Gene orthology analysis

Orthologue identification was performed using OrthoFinder version 1.1.10 (Emms and Kelly, 2015) on predicted proteins from all sequenced *P. fragariae* and *P. rubi* isolates. Orthogroups were investigated for expanded and unique groups for races UK1, UK2 and UK3. Unique orthogroups were those containing proteins from only one race and expanded orthogroups were those containing more proteins from a specific race than other races. Venn diagrams were created with the VennDiagram R package version 1.6.20 (Chen and Boutros, 2011) in R version 3.2.5 (R Core Team, 2016).

### Identification and analysis of variant sites and population structure

Variant sites were identified through the use of the GATK version 3.6 HaplotypeCaller in diploid mode (McKenna et al., 2010) following alignment of Illumina reads for all sequenced isolates to the FALCON-Unzip assembled genome with Bowtie 2 version 2.2.6 (Langmead and Salzberg, 2012) and filtered with vcflib (Garrison, 2012) and VCFTools (Danecek et al., 2011). Additionally, structural variants were identified with SvABA (Wala et al., 2018) following the alignment of Illumina reads from all sequenced isolates to the FALCON-Unzip assembled genome with BWA-mem version 0.7.15 (Li, 2013). Population structure was assessed with fastSTRUCTURE (Raj et al., 2014) following conversion of the input file with Plink version 1.90 beta (Purcell et al., 2007). Finally, a custom Python script was used to identify variant sites private to races UK1, UK2 or UK3 (see Availability of Computer Code below).

### Assessment of gene expression levels and the identification of candidate avirulence genes

RNA-Seq reads of BC-1, BC-16 and NOV-9 were aligned to the assembled genomes of BC-1, BC-16 and NOV-9 as described above. Expression levels and differentially expressed genes were identified with featureCounts version 1.5.2 (Liao et al., 2014) and the DESeq2 version 1.10.1 R package (Love et al., 2014). Multiple test correction was performed as part of the analysis within the DESeq2 package.

Candidate avirulence genes were identified through the analysis of uniquely expressed genes and uniquely differentially expressed genes for each of the three isolates. Uniquely expressed genes were those with a Fragments Per Kilobase of transcript per Million mapped reads (FPKM) value greater than or equal to 5 in any time point from the inoculation time course experiment for a single isolate. Uniquely differentially expressed genes were those with a minimum LFC of 3 and a *p*-value threshold of 0.05. Genes were compared between isolates through the use of orthology group assignments described above and scored on a one to six scale, with five and six deemed high confidence and one and two deemed as low confidence.

To identify homologous genes in other *Phytophthora* spp., candidate RxLRs were processed by SignalP-5.0 (Armenteros et al., 2019) to identify the cleavage site. The signal peptide sequence was removed before being submitted for a tblastn (Altschul et al., 1997) search on GenBank; interesting top hits are reported.

The expression levels of a strong candidate *PfAvr2* gene, PF003_g27513 and a candidate *PfAvr3* gene, PF009_g26276, were assessed via RT-qPCR. RNA-Seq results for reference genes identified previously in *Phytophthora parasitica* (Yan and Liou, 2006) were examined for stability of expression levels, resulting in the selection of β-tubulin (PF003_g4288) and WS41 (PF003_g28439) as reference genes. Primers were designed using the modified Primer3 version 2.3.7 implemented in Geneious R10 (Untergasser et al., 2012). Reverse transcription was performed on three biological replicates of each sequenced timepoint, alongside: 24 hpi, 72 hpi and 96 hpi for BC-1 and 24 hpi, 48 hpi and 96 hpi for NOV-9 with the QuantiTect Reverse Transcription Kit. Quantitative PCR (qPCR) was then performed in a CFX96^TM^ Real-Time PCR detection system in 10 µL reactions of: 5 µL of 2x qPCRBIO SyGreen Mix Lo-Rox, 2 µL of a 1:3 dilution of the cDNA sample in dH_2_O and 0.4 µL of each 10 µM primer and 2.2 µL dH_2_O. The reaction was run with the following conditions: 95 °C for 3 minutes, 39 cycles of 95 °C for 10 seconds, 62 °C for 10 seconds and 72 °C for 30 seconds. This was followed by 95 °C for 10 seconds, and a 5 second step ranging from 65 °C to 95 °C by 0.5 °C every cycle. At least two technical replicates for each sample were performed and the melt curve results were analysed to ensure the correct product was detected. Relative gene expression was calculated using the comparative cycle threshold (C_T_) method (Livak and Schmittgen, 2001). Where there was a difference of at least 1 CT value between the minimum and maximum results for technical replicates, further reactions were conducted. Outlier technical replicates were first identified as being outside 1.5 times the interquartile range. Following this, additional outliers were identified using the Grubb’s test and excluded from the analysis. Technical replicates were averaged for each biological replicate and expression values were calculated as the mean of the three biological replicates.

### Investigation of *cis* and *trans* variations for strong candidate avirulence gene

The regions upstream and downstream of PF003_g27513 and PF009_g26276 in the FALCON-Unzip assembly of BC-16 and the SMARTdenovo assembly of NOV-9 were aligned with MAFFT in Geneious R10 (Katoh et al., 2002; Katoh and Standley, 2013). Variant sites were identified through visual inspection of the alignment alongside visualisation of aligned short reads from BC-16 and NOV-9 to both assemblies with Bowtie 2 version 2.2.6 (Langmead and Salzberg, 2012) in IGV (Thorvaldsdóttir et al., 2013).

Transcription factors were identified with a previously described HMM of transcription factors and transcriptional regulators in Stramenopiles (Buitrago-Flórez et al., 2014). These proteins were then analysed through methods described above for gene loss or gain, nearby variant sites and differential expression between isolates.

### Availability of computer code

All computer code is available at: https://github.com/harrisonlab/phytophthora_fragariae,

https://github.com/harrisonlab/phytophthora_rubi and https://github.com/harrisonlab/popgen/blob/master/snp/vcf_find_difference_pop.py.

## RESULTS

### Race typing allowed standardisation of US and UK race nomenclature

The *P. fragariae* isolates A4 (race US4), BC-1 (race CA1), BC-16 (race CA3), NOV-5 (race CA1), NOV-9 (race CA2), NOV-27 (race CA2) and NOV-71 (race CA2) (**Table 1**) were phenotyped on a differential series of four *F.* × *ananassa* cultivars with known resistance. These were: ‘Allstar’ (containing *Rpf1*, *Rpf2* and *Rpf3*), ‘Cambridge Vigour’ (containing *Rpf2* and *Rpf3*), ‘Hapil’ (containing no resistance genes) and ‘Redgauntlet’ (containing *Rpf2*) (van de Weg et al., 1996; R. Harrison, unpublished). Following assessment of below ground symptoms, it was shown that race CA1 was equivalent to UK1, race CA2 was equivalent to UK3, race CA3 was equivalent to UK2 and race US4 was equivalent to UK2 (**Figure 1**; **Table 2**).

**FIGURE 1.**
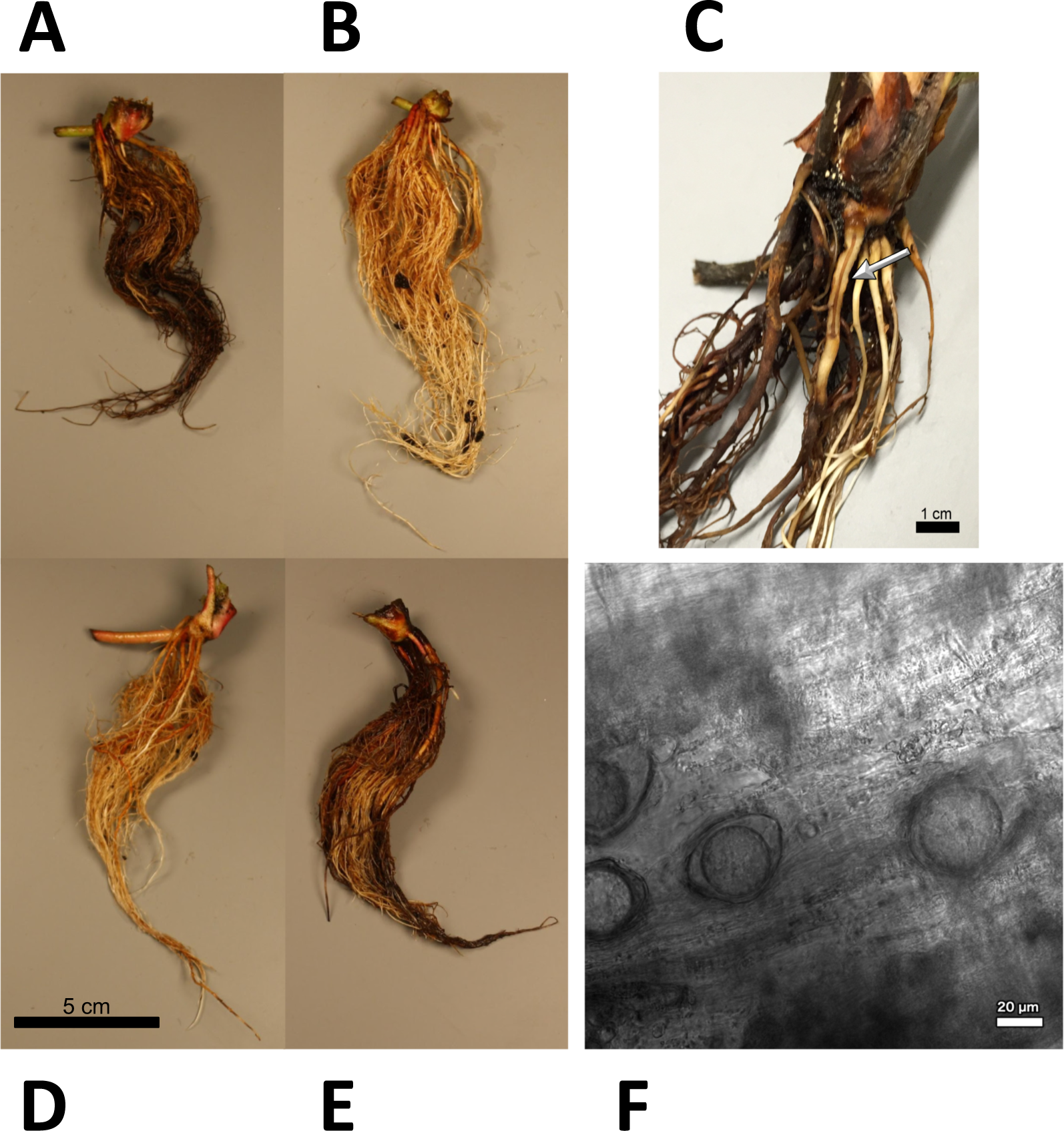
Observed *Phytophthora fragariae* symptoms in the cultivated strawberry (*Fragaria* × *ananassa*). **(A,B,D,E)** Roots of *Fragaria* × *ananassa* harvested six weeks after inoculation with *Phytophthora fragariae* mycelial slurry. **(A)** Successful infection of the *P. fragariae* isolate NOV-27 (race CA2) on a susceptible ‘Redgauntlet’ plant (*Rpf2* only). **(B)** Unsuccessful infection of the *P. fragariae* isolate A4 (US4/UK2) on a resistant ‘Redgauntlet’ plant (*Rpf2* only). **(C)** Example of “red core” symptoms observed in ‘Hapil’ roots infected with BC-16, three weeks post inoculation. **(D)** Unsuccessful infection of the *P. fragariae* isolate NOV-27 (race CA2) on a resistant ‘Allstar’ plant (*Rpf1*, *Rpf2* and *Rpf3*). **(E)** Successful infection of the *P. fragariae* isolate A4 (race US4/UK2) on a susceptible ‘Hapil’ plant (no *Rpf* genes). **(F)** Example of BC-16 oospores observed in ‘Hapil’ roots, 3 weeks post inoculation, confirming infection.

**TABLE 2.**
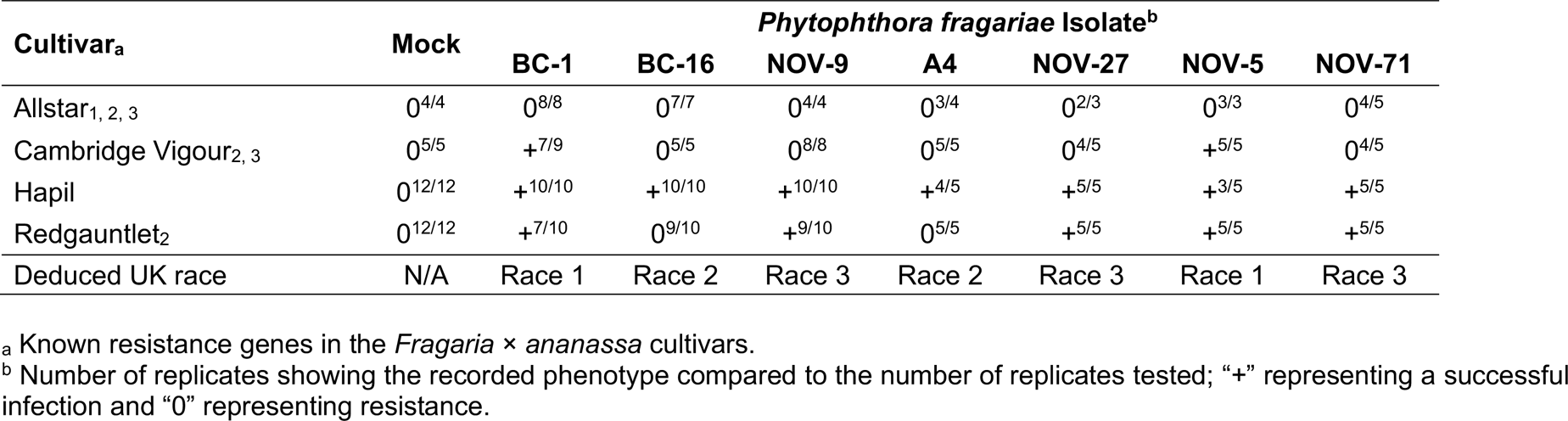
| **Phytophthora fragariae race structure detected by three differential strawberry (Fragaria × ananassa) accessions and categorisation into UK race scheme.**

### A highly contiguous genome assembly of BC-16

Long read PacBio sequencing generated a highly contiguous *P. fragariae* isolate BC-16 (UK2) reference genome of 91 Mb in 180 contigs with an N50 value of 923.5 kb (**Table 3**). This was slightly larger than the closely related, well studied species from Clade 7, *P. sojae*, at 83 Mb (**Table 3**; Tyler et al., 2006). The BC-16 assembly contained 266 of 303 eukaryotic Benchmarking Universal Single-Copy Orthologs (BUSCO) genes (**Table 4**), compared to 270 in *P. sojae* (Armitage et al., 2018) and so likely represented a similar completeness of the genome as the *P. sojae* assembly. The *P. fragariae* genome was shown to be highly repeat rich, with 38% of the assembly identified as repetitive or low complexity, a larger value than the 29% shown for *P. sojae* (Armitage et al., 2018). A total of 37,049 genes encoding 37,346 proteins were predicted in the BC-16 assembly, consistening of 20,222 genes predicted by BRAKER1 (Hoff et al., 2016) and 17,131 additional genes added from CodingQuarry (Testa et al., 2015). From these gene models, 486 putative RxLR effectors, 82 putatitve crinkler effectors (CRNs) and 1,274 putative apoplastic effectors were identified (**Table 5**). Additionally, a total of 4,054 low confidence gene models were added from intergenic ORFs identified as putative effectors. These consisted of 566 putative RxLRs, 5 putative CRNs and 3,483 putatitve apoplastic effectors. This resulted in a combined total of 41,103 genes encoding 41,400 proteins with 1,052 putative RxLRs, 85 putative CRNs and 4,757 putative apoplastic effectors (**Table 3** and **Table 5**).

**TABLE 3.** | Long read PacBio sequencing generated a highly contiguous *Phytophthora fragariae* BC-16 (UK2) reference genome. Assembly and annotation statistics compared to *Phytophthora sojae* P6497 (Tyler et al., 2006). Values in brackets include low confidence gene models from open reading frames.

|  | *Phytophthora fragariae*<br>**BC-16** | *Phytophthora sojae*<br>**P6497** |
| --- | --- | --- |
| Assembly size (Mb) | 90.97 | 82.60 |
| Number of contigs | 180 | 862 |
| N50 (kb) | 923.5 | 386.0 |
| L50 | 33 | 61 |
| Repeatmasked | 38% | 29% |
| Genes | 37,049 (41,103) | 26,584 |
| RxLRs | 486 (1,052) | 350 |
| CRNs | 82 (85) | 40 |
| Apoplastic effectors | 1,274 (4,757) | N/A |

**TABLE 4.**
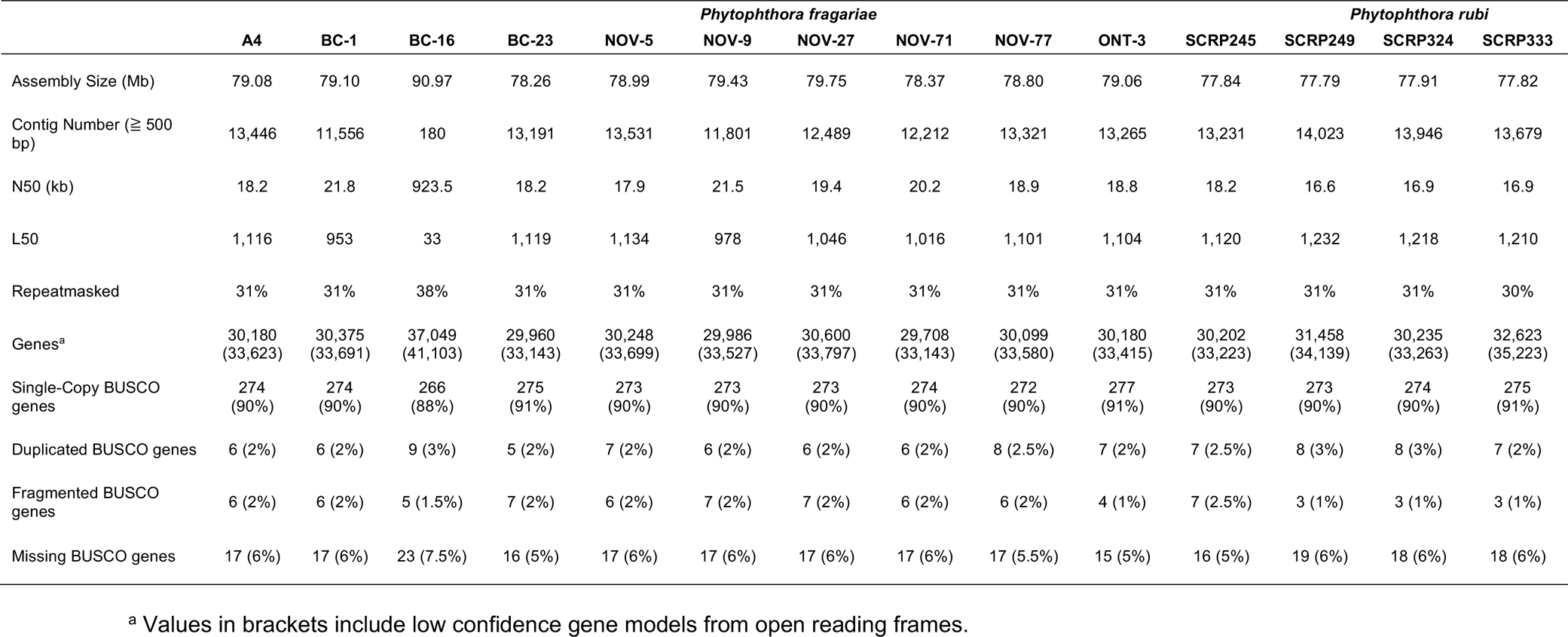
| **Comparable assembly statistics and gene predictions in resequenced isolates of Phytophthora fragariae and Phytophthora rubi.**

**TABLE 5.** | **Effector gene predictions in resequenced isolates of *Phytophthora fragariae* and *Phytophthora rubi*.**

|  | Phytophthora fragariae |  |  |  |  |  |  |  |  |  | Phytophthora rubi |  |  |  |
| --- | --- | --- | --- | --- | --- | --- | --- | --- | --- | --- | --- | --- | --- | --- |
|  | A4 | BC-1 | BC-16 | BC-23 | NOV-5 | NOV-9 | NOV-27 | NOV-71 | NOV-77 | ONT-3 | SCR245 | SCR249 | SCR324 | SCR333 |
| Secreted proteins<sup>a</sup> | 3,637 | 3,601 | 4,217 | 3,611 | 3,626 | 3,637 | 3,690 | 3,633 | 3,620 | 3,658 | 3,581 | 3,697 | 3,683 | 3,832 |
| RxLR HMM<sup>b</sup> | 194 | 194 | 218 | 188 | 196 | 186 | 183 | 191 | 166 | 191 | 196 | 197 | 195 | 207 |
| RxLR-EER Regex<sup>c</sup> | 178 | 184 | 208 | 176 | 186 | 174 | 172 | 182 | 150 | 178 | 185 | 188 | 176 | 188 |
| RxLR Regex | 371 | 367 | 445 | 364 | 370 | 356 | 369 | 374 | 341 | 367 | 370 | 363 | 350 | 359 |
| Final RxLR effectors<sup>d</sup> | 410<br>(935) | 405<br>(919) | 486<br>(1,052) | 402<br>(928) | 408<br>(918) | 397<br>(951) | 405<br>(899) | 412<br>(931) | 378<br>(934) | 403<br>(898) | 407<br>(882) | 407<br>(882) | 395<br>(861) | 410<br>(826) |
| CRN LFLAK HMM<sup>c</sup> | 85 | 83 | 114 | 86 | 84 | 82 | 87 | 90 | 86 | 90 | 90 | 156 | 93 | 154 |
| CRN DWL HMM<sup>c</sup> | 95 | 87 | 121 | 96 | 91 | 96 | 93 | 101 | 83 | 90 | 87 | 139 | 102 | 142 |
| Final CRNs<sup>d,e</sup> | 55<br>(68) | 53<br>(59) | 82<br>(87) | 55<br>(62) | 53<br>(67) | 50<br>(61) | 57<br>(63) | 59<br>(71) | 62<br>(75) | 61<br>(74) | 60<br>(68) | 128<br>(132) | 71<br>(85) | 128<br>(134) |
| Apoplatic effectors<sup>d,f</sup> | 991<br>(3,896) | 1,002<br>(3,798) | 1,274<br>(4,757) | 980<br>(3,630) | 986<br>(3,913) | 1,007<br>(3,983) | 1,011<br>(3,708) | 1,007<br>(3,911) | 1,010<br>(3,922) | 982<br>(3,709) | 984<br>(3,522) | 1,017<br>(3,266) | 1,059<br>(3,607) | 1,090<br>(3,268) |

<sup>a</sup> SignalP: Nielsen et al., 1997; Bendtsen et al., 2004; Petersen et al., 2011 and Phobius: Käll et al., 2007.
<sup>b</sup> Whisson et al., 2007.
<sup>c</sup> Armitage et al., 2018.
<sup>d</sup> Values in brackets include low confidence gene models from open reading frames.
<sup>e</sup> Both LFLAK and DWL HMM models present (Armitage et al., 2018).
<sup>f</sup> ApoplastP: Spersneider et al., 2018.

### A comparable number of effector genes were identified through resequencing of isolates of ***Phytophthora fragariae* and *Phytophthora rubi***

Ten isolates of *P. fragariae* and three isolates of *P. rubi* were additionally resequenced, *de novo* assembled and annotated (**Table 1** and **Table 4**). These isolates showed similar assembly statistics within and between species; an average of 79 Mb in 12,804 contigs with an N50 of 19.3 kb in *P. fragariae*, compared to an average of 78 Mb in 13,882 contigs with an N50 of 16.8 kb in *P. rubi*. All assemblies showed 31% of the assembly was identified as repetitive or low complexity sequences. On average both *P. fragariae* and *P. rubi* showed high levels of completeness, with an average of 274/303 BUSCO genes identified as single copies in these assemblies. Interestingly, a total of 17 genes were consistently not identified in Illumina, PacBio and Nanopore assemblies of *P. fragariae* and *P. rubi* isolates, suggesting these genes may be absent from these species and as such may not be considered true eukaryotic BUSCO genes. An average of 67 CRNs were predicted in *P. fragariae* and an average of 118 CRNs in *P. rubi*. A significantly (*p* = 0.013) smaller number of RxLRs were predicted in *P. rubi* isolates than *P. fragariae* isolates alongside a significantly (*p* = 0.039) smaller number of apoplastic effectors predicted in *P. rubi* than *P. fragariae* (**Table 5**). However, it is important to note that these effectors were predicted from a subset of secreted proteins, of which there were significantly (*p* = 0.030) fewer predicted in *P. rubi* than in *P. fragariae* (**Table 5**).

### No distinguishing gene loss or gain; or the presence of INDELs or SNPs in candidate avirulence genes could be found associated with race variation

An orthology analysis of all predicted proteins from the isolates of *P. fragariae* and *P. rubi* assigned 481,942 (98.7%) proteins to 38,891 orthogroups. Of these groups, 17,101 contained at least one protein from all fourteen sequenced isolates and 13,132 of these groups consisted entirely of single-copy proteins from each isolate. There were 2,345 of these groups unique to *P. rubi* and 1,911 of these groups unique to *P. fragariae*. Analysis of unique and expanded orthogroups for isolates of the UK1, UK2 and UK3 races did not lead to the identification of candidate avirulence genes, as these groups did not contain putative effector genes (**Figure 2**).

**FIGURE 2.**
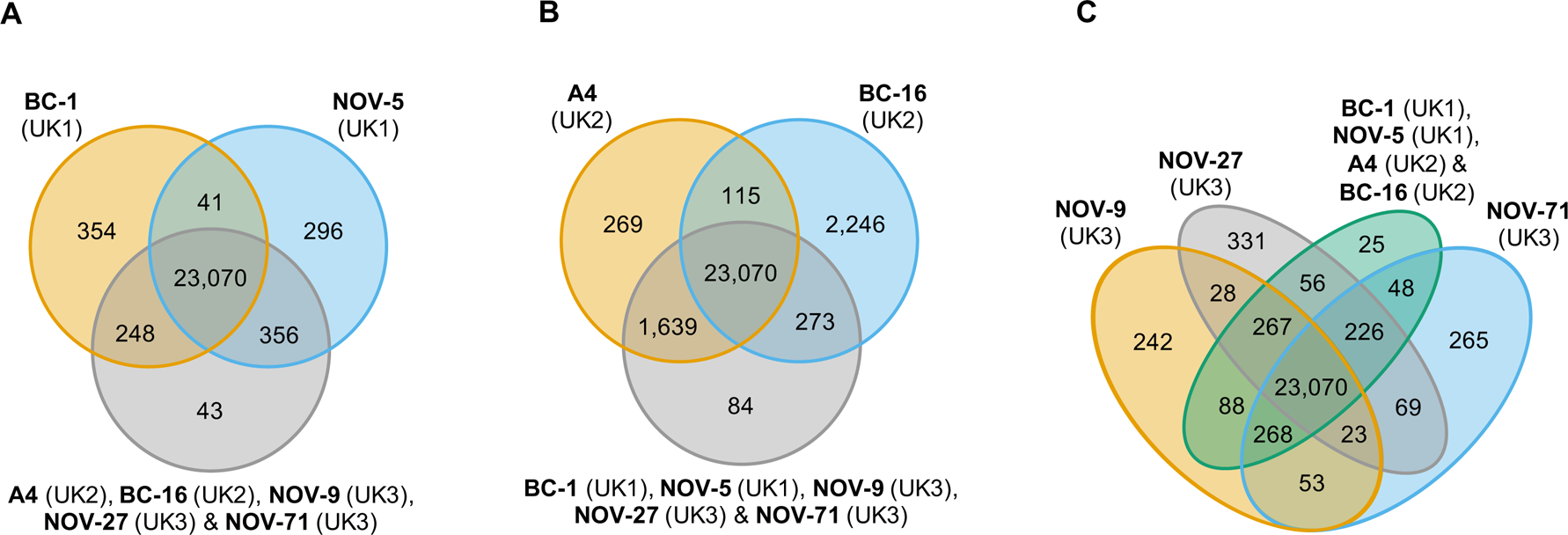
Analysis of unique and expanded orthogroups for *Phytophthora fragariae* isolates of the UK1, UK2 and UK3 races did not lead to the identification of candidate avirulence genes. Orthology groups were identified by OrthoFinder (Emms and Kelly, 2015) and Venn diagrams were plotted using the VennDiagram R package (Chen and Boutros, 2011) in R (R Core Team, 2016). **(A)** Analysis focused on the *P. fragariae* isolates of race UK1: BC-1 and NOV-5 compared to isolates of races UK2 and UK3. **(B)** Analysis focused on the *P. fragariae* isolates of race UK2: A4 and BC-16 compared to isolates of races UK1 and UK3. **(C)** Analysis focused on the *P. fragariae* isolates of race UK3: NOV-5, NOV-27 and NOV-71 compared to isolates of races UK1 and UK2.

A total of 725,444 SNP sites and 95,478 small INDELs were identified within the *P. fragariae* and *P. rubi* isolates by the Genome Analysis ToolKit (GATK) haplotypecaller (McKenna et al., 2010) and 80,388 indels and 7,020 structural variants were identified by SvABA (Wala et al., 2018). Analysis of high quality, biallelic SNP sites allowed the identification of a distinct population consisting of the isolates of race UK1, UK2 and UK3, hereafter referred to as the UK1-2-3 population, with SCRP245, the only UK isolate, potentially forming an ancestral or hybrid isolate between the UK1-2-3 population and the population represented by BC-23 and ONT-3 (**Figure 3**; Supplementary Figure S2). Additionally, clear separation between isolates of *P. fragariae* and *P. rubi* was observed. Further analysis within the UK1-2-3 population allowed for the identification of private variants, which were only present in isolates of one of the three races in this population. This resulted in the identification of eleven private variants in the UK2 race shared between both A4 and BC-16; however, neither the genes they fell within or were neighbours to were predicted to encode effectors or secreted proteins and likely did not explain the differences in pathogenicity.

**FIGURE 3.**
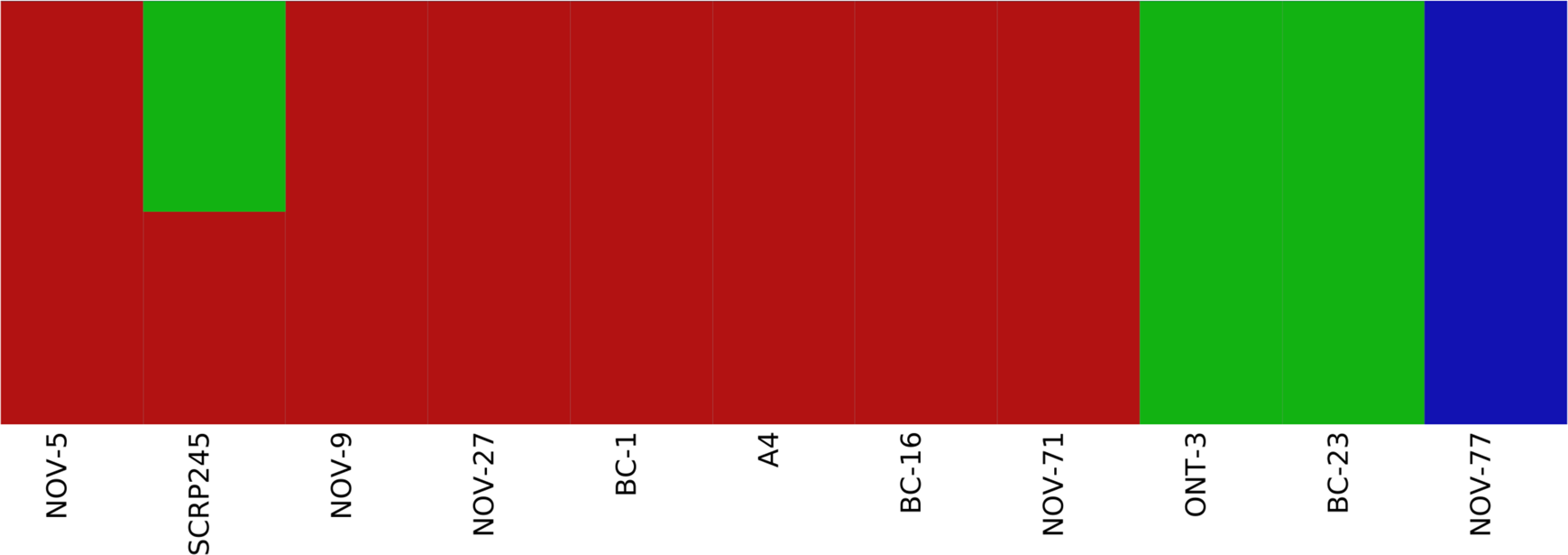
Analysis of high quality, biallelic SNP sites split the isolates into three populations with SCRP245, potentially forming an ancestral or hybrid isolate between the UK1-2-3 population and the population represented by BC-23 and ONT-3. Distruct plot of fastSTRUCTURE (Raj et al., 2014) results carried out on all sequenced isolates of *Phytophthora fragariae*. Each colour represents a different population. Variant sites were identified by aligning Illumina reads of all the sequenced isolates to the reference assembly of the BC-16 isolate of *P. fragariae* with Bowtie 2 (Langmead and Salzberg, 2012) and analysis with the Genome Analysis Toolkit (GATK) haplotypecaller (McKenna et al., 2010). Sites were filtered with VCFtools (Danecek et al., 2011) and VCFlib (Garrison, 2012) to leave only high quality, biallelic SNP sites.

### Wide scale transcriptional reprogramming of effectors during strawberry infection

RNA-Seq data were generated from an inoculation time course experiment for representatives of races UK1, UK2 and UK3; BC-1, BC-16 and NOV-9 respectively. Following determination of expression levels of the predicted genes in both *in planta* and mycelia samples, a correlation analysis showed biological replicates of each timepoint grouped together (**Figure 4**). Additionally, the 48 hours post inoculation (hpi) BC-1 time point grouped with the BC-16 24 hpi time point, suggesting these may represent similar points in the infection process. However, clear separation between the BC-16 timepoints was observed (**Figure 4**). A total of 13,240 transcripts from BC-16 (32%) showed expression above a FPKM value threshold of five in at least one sequenced BC-16 time point and 9,329 (23%) of these showed evidence of differential expression in at least one sequenced *in planta* time point compared to mycelium grown in artificial media, suggesting large scale transcriptional reprogramming. As three time points post inoculation were sequenced for BC-16, the changes in expression over the course of the infection process was investigated. A total of 2,321 transcripts (6%) showed a Log2 Fold Change (LFC) greater than or equal to one or less than or equal to minus one, representing a general reprofiling during infection in comparison to growth in artificial media. Of transcripts differentially expressed *in planta* compared to artificial media, fewer transcripts were differentially expressed at both 24 hpi and 96 hpi than between sequential timepoints (297 compared to 1,016 and 1,604 transcripts). This suggested that changes during the progress of infection were captured by this dataset (**Figure 5**).

**FIGURE 4.**
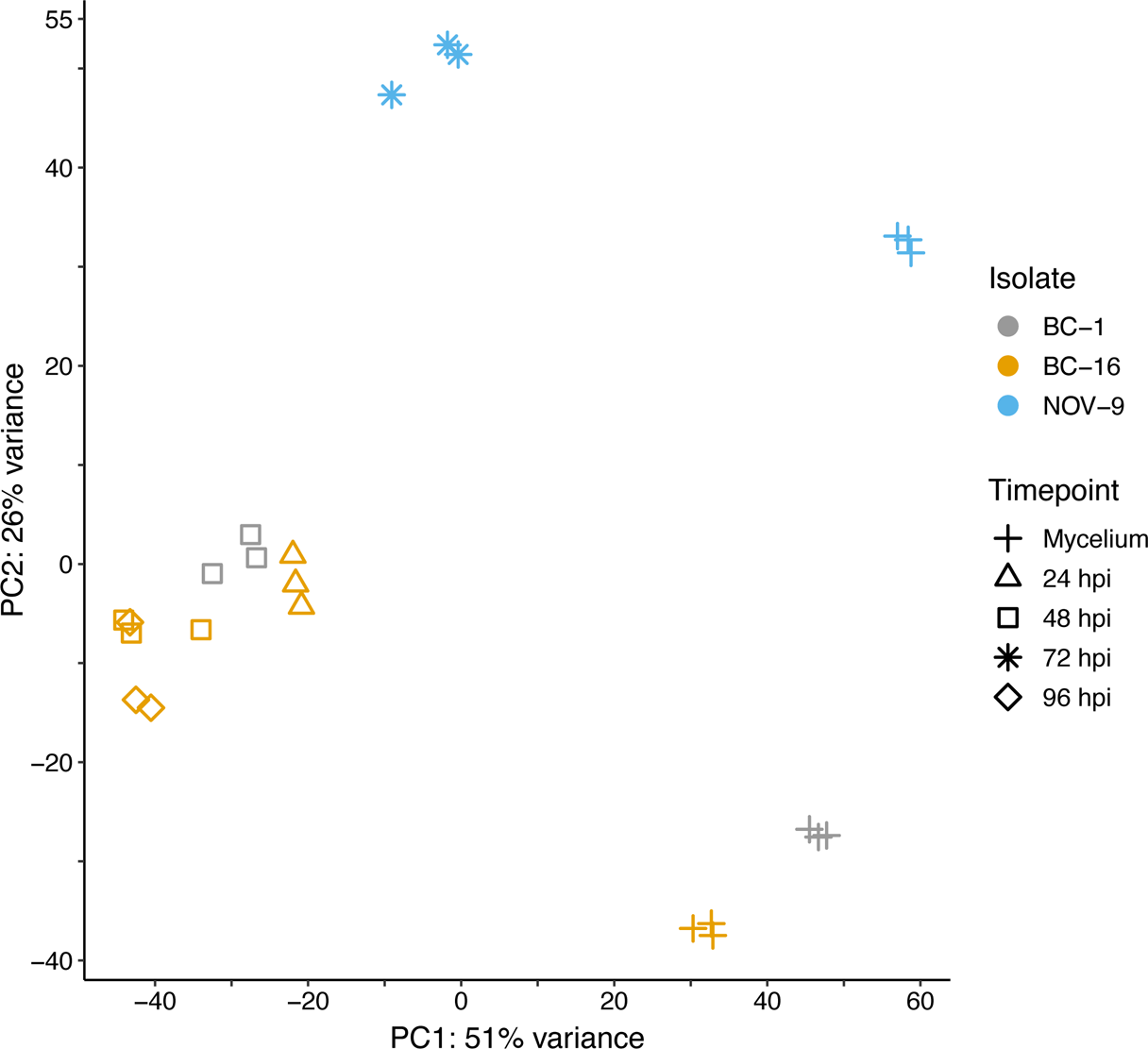
Principal component analysis of expression data showed biological replicates of each timepoint grouped together, with clear separation observed for the BC-16 timepoints. RNA-Seq reads were aligned to the assembly of the BC-16 isolate of *Phytophthora fragariae* using STAR version 2.5.3a (Dobin et al., 2013). Predicted transcripts were then quantified with featureCounts version 1.5.2 (Liao et al., 2014) and differential expression was identified with the DESeq2 version 1.10.1 R package (Love et al., 2014). Following this, an rlog transformation of the expression data was plotted as a principal component analysis with R (R Core Team, 2016).

**FIGURE 5.**
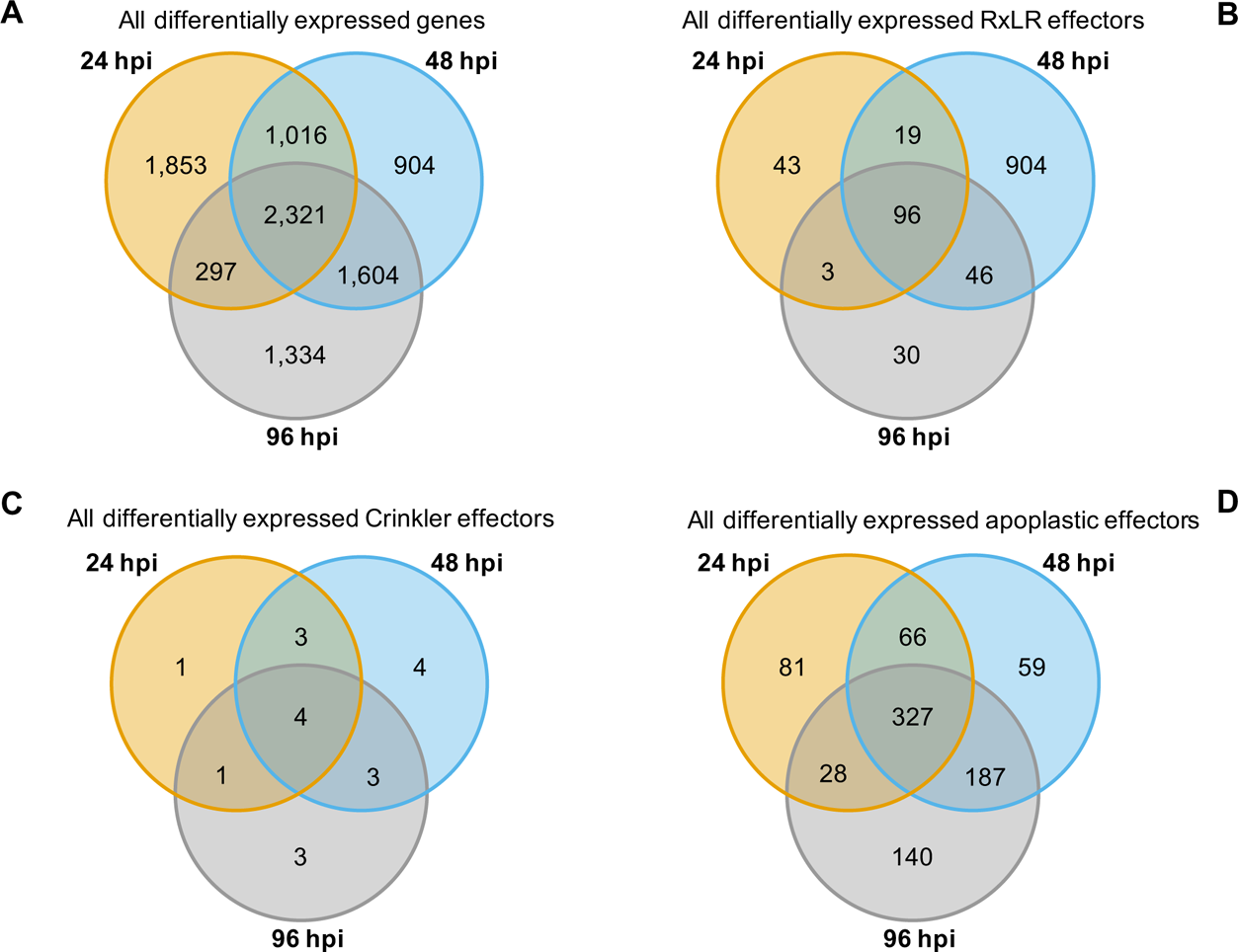
*In planta* RNA-Seq dataset captured changes during the progress of infection in the *Phytophthora fragariae* BC-16 isolate. RNA-Seq reads were aligned to the assembly of the BC-16 isolate of *P. fragariae* using STAR (Dobin et al., 2013). Predicted transcripts were then quantified with featureCounts (Liao et al., 2014) and differential expression was identified with DESeq2 (Love et al., 2014). Following this, Venn diagrams were plotted using the VennDiagram R package (Chen and Boutros, 2011) in R (R Core Team, 2016). **(A)** All differentially expressed genes. **(B)** All differentially expressed RxLR effectors. **(C)** All differentially expressed Crinkler effectors. **(D)** All differentially expressed putative apoplastic effectors.

Levels of expression of effector genes were also assessed. A total of 274 (26%) RxLR effectors, 27 (31%) CRNs and 880 (18%) putative apoplastic effectors from BC-16 showed evidence of expression above the FPKM threshold of 5 in at least one sequenced time point. The majority of these effector genes also showed evidence of differential expression during *in planta* time points compared to *in vitro* mycelium. A total of 253 (24%) RxLR effectors, 19 (22%) CRNs and 888 (18%) putative apoplastic effectors showed an LFC greater than or equal to one or less than or equal to minus one, representing wide scale transcriptional reprogramming of effector genes during infection (**Figure 5**). Ranking of the 50 highest expressed genes *in planta* with a LFC of ≥3 in comparison to its respective mycelium, identified four putative RxLR genes that were upregulated by all three isolates (**Table 6**). These genes represent putative core *P. fragariae* RxLR’s important for pathogenicity on strawberry. Interestingly, a BLASTP search of one of these putative core effectors, PF003_g16448 (amino acids 19-139), showed homology to *P. sojae Avr1b* (58 % pairwise identity), GenBank accession AF449625 (Shan et al., 2004).

**TABLE 6.** | **Conserved putative core Phytophthora fragariae RxLR effector candidates important for pathogenicity on strawberry (Fragaria × ananassa) from isolates BC-1, BC-16 and NOV-9.** Conserved RxLR candidates in the 50 highest expressed genes *in planta* with a log fold change (LFC) of ≥3, based on comparison to respective mycelium. The BC-16 24 hour post inoculation (hpi) timepoint was used for BC-16.

| Orthogroup | BC-16 gene ID | Fragments Per Kilobase of transcript per Million mapped reads (FPKM) |  |  |  |  |  |  |  | Avr homology |
| --- | --- | --- | --- | --- | --- | --- | --- | --- | --- | --- |
|  |  | BC-1 (UK1) |  | BC-16 (UK2) |  |  |  | NOV-9 (UK3) |  |  |
|  |  | Mycelium | 48 hpi | Mycelium | 24 hpi | 48 hpi | 96 hpi | Mycelium | 72 hpi |  |
| OG0010423 | PF003_g35418 | 94 | 5,354 | 192 | 6,578 | 4,986 | 2,961 | 12 | 3,868 |  |
| OG0018019 | PF003_g6480 | 241 | 3,544 | 235 | 2,113 | 1,977 | 1,717 | 231 | 2,501 |  |
| OG0011458 | PF003_g16448 | 4 | 2,899 | 4 | 4,656 | 2,108 | 1,090 | 7 | 1,699 | *P. sojae Avr1b* |
| OG0021012 | PF003_g16234 | 36 | 1,345 | 31 | 1,181 | 827 | 674 | 23 | 737 |  |

<sup>a</sup> Top hit from GenBank tblastn search is AF449623, *Phytophthora sojae Avr1b* (Shan et al., 2004).

### RxLR effector PF003_g27513 is a strong candidate for *PfAvr2*

Comparing transcripts with the highest LFC (*in planta* vs mycelium) between isolates led to the identification of the putative RxLR effector encoding transcript PF003_g27513.t1 as a potential candidate for *PfAvr2* in BC-16. PF003_g27513 had a peak FPKM value in BC-16 of 9,392 compared to peaks of 5 and 34, in BC-1 and NOV-9, respectively (Supplementary Table S2). Subsequent RT-qPCR analysis of further timepoints in the *in planta* timecourse supported the findings of the RNA-Seq timepoints and showed the expression of putative *PfAvr2* in the UK2 isolates BC-16 and A4 was significantly (*p* < 0.05) different to those from all samples of BC-1 (UK1) and NOV-9 (UK3; **Figure 6**). A BLASTP search of PF003_g16448 (amino acids 26-137) revealed homology to *P. sojae Avh6/Avr1d* (37 % pairwise identity), GenBank accession JN253642 (Wang et al., 2011; Na et al., 2013).

**FIGURE 6.**
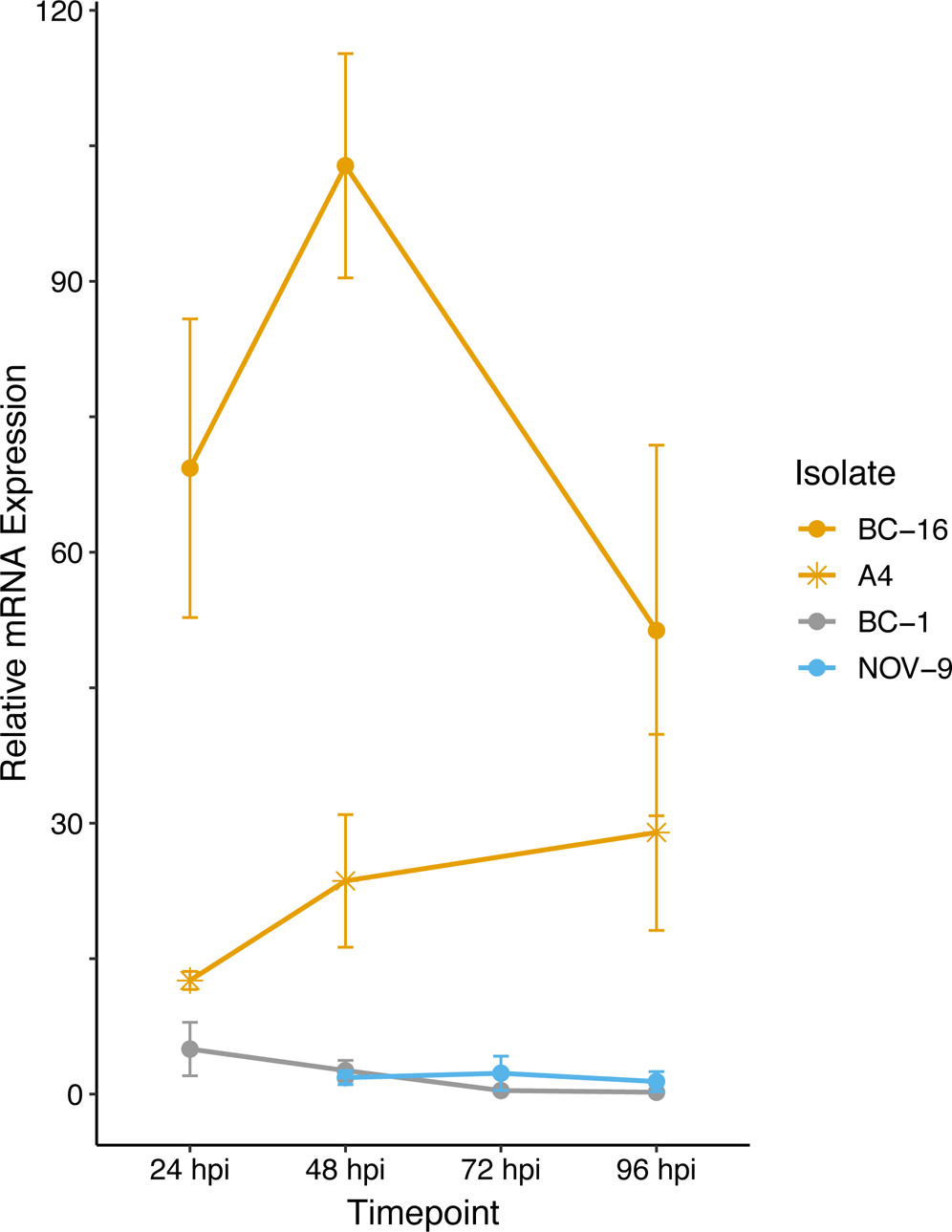
*PfAvr2* candidate PF003_g27513 is differentially expressed in *Phytophthora fragariae* UK-1-2-3 isolates. Quantitative reverse transcription PCR of a strong candidate for the avirulence gene possessed by BC-16 and A4, but not BC-1 and NOV-9 (PF003_g27513.t1). Plots created by the ggplot2 R package (Wickham, 2016) in R version 3.4.3 (R core team, 2017).

The surrounding sequence of putative *PfAvr2* in BC-16, BC-1 and NOV-9 was investigated for sequence variants that may explain the expression difference. As no variants were identified near the gene from the above mentioned variant panel, an assembly of the NOV-9 isolate was created from Nanopore sequencing data, producing an assembly of 93.72 Mbp in 124 contigs with an N50 of 1,260 Kb. This resulted in the identification of a SNP from T in BC-16 to G in NOV-9 ∼14 Kb downstream of the stop codon and a 30 bp insertion in NOV-9 ∼19 Kb upstream of the start codon (**Figure 7A**). The upstream variant appeared to be a sequencing error following the investigation of the alignment of short reads of BC-16 and NOV-9 to each assembly and so was rejected. Expression of the genes surrounding PF003_g27513 was investigated and PF003_g27514, directly upstream of putative *PfAvr2* is expressed by all three isolates (Supplementary Table S2).

**FIGURE 7.**
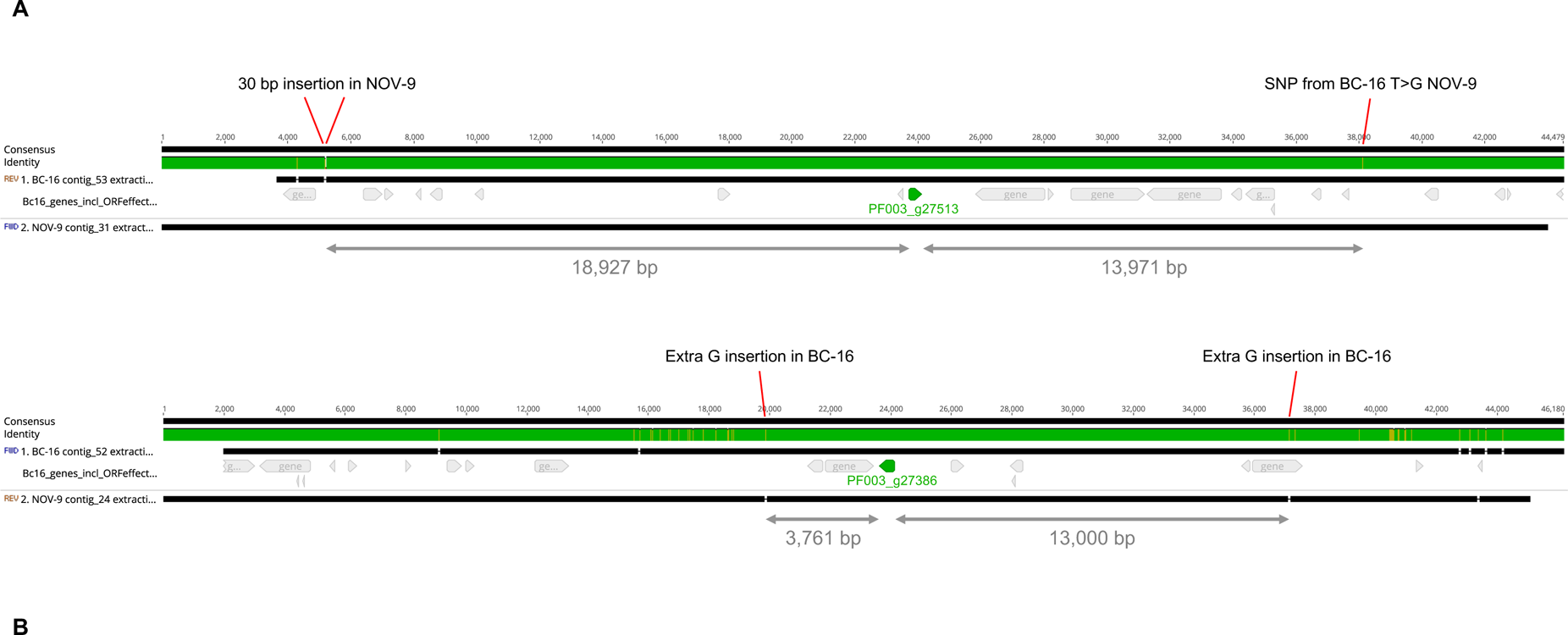
Differential expression of putative *PfAvr2* and *PfAvr3* is not due to sequence variation in *Phytophthora fragariae* BC-16 (UK2) and NOV-9 (UK3) genomes. Regions surrounding candidate avirulence genes, *PfAvr2* and *PfAvr3*, from BC-16 and NOV-9 aligned with MAFFT in Geneious R10 (Katoh et al., 2002; Katoh and Standley, 2013). (**A**) Putative *PfAvr2*, showing a 30 bp insertion in the NOV-9 sequence upstream of PF003_g27513 and a T to G SNP in NOV-9 downstream of the gene of interest. (**B**) Putative *PfAvr3*, showing an extra G insertion in BC-16 upstream of PF003_g27386 (an orthologue of PF009_g26267) and an extra G insertion in BC-16 3,761 bp downstream.

Additionally, putative transcription factors and transcriptional regulators were identified in all sequenced genomes. This resulted in the identification of 269 genes in the BC-16 isolate of *P. fragariae*. However, analysis of gene loss or gain and an investigation of variant sites showed no race specific differences, though expression level variation was observed.

### RxLR effector PF009_g26276 is a putative candidate for *PfAvr3*

Further analysis of the RNA-Seq datasets identified the putative RxLR effector encoding transcript PF009_g26276 (an orthologue of PF003_g27386) as a potential candidate for *PfAvr3* in NOV-9, with a peak FPKM value in NOV-9 of 199 compared to *in planta* peaks of 6 and 12 in BC-1 and BC-16, respectively (Supplementary Table S3). Similar to putative *PfAvr2*, no sequence differences in putative *PfAvr3* were observed between the three isolates. Analysis of the surrounding region between NOV-9 and BC-16 showed no sequence differences 13,000 bp upstream and 3,761 bp downstream of PF009_g26276 (**Figure 7B**). The two genes upstream from putative *PfAvr3* are not expressed in any isolate (Supplementary Table S3), whereas the gene directly downstream in BC-16, PF003_g27385, is expressed by all the three isolates analysed.

Subsequent RT-qPCR analysis of further timepoints in the *in planta* time course revealed that the putative *PfAvr3* was not expressed during any timepoints by the UK1 isolate BC-1 or by the UK2 isolates BC-16 and A4 (Supplementary Figure S3). However, absolute expression of PF009_g26276 in NOV-9 remained low. Expression in NOV-9 at 72 hpi was significantly (*p* < 0.05) greater than those from all samples of BC-1, BC-16 and A4.

### Candidate genes for *PfAvr1* identified from expression level variation

Analysis of the RNA-Seq datasets did not reveal an obvious candidate for *PfAvr1*. Further analysis using custom scripts identified genes, which were uniquely expressed in BC-1 and those uniquely differentially expressed. These genes were scored for confidence of race avirulence determinant; high, medium and low. No genes were identified in the high confidence class in BC-1, but four genes were scored as medium confidence, one of which was a putative apoplastic effector (**Supporting Candidate Table**). The analysis was repeated for BC-16 and NOV-9, in case the putative RxLR candidates are not *PfAvr*2 and *PfAvr3*, respectively (**Supporting Candidate Table**).

## DISCUSSION

Understanding pathogenicity of plant pathogens is critical for developing durable resistance strategies. *P. fragariae* is a continuing threat to strawberry production. The UK1-2-3 population displayed clear separation from the other isolates of *P. fragariae* in this study. Two putative avirulence candidates for UK2 and UK3 were identified through population resequencing and analyses of gene expression during *P. fragariae* infection. We have shown that there are no distinguishing gene loss or gain events, INDELs or SNPs associated with race variation in UK1-2-3. Our results also suggest that the polymorphisms associated with avirulence to race UK2 and UK3 resistance is controlled *in trans* or with other stable forms of epigenetic modulating gene expression.

This study utilised long read sequencing technologies to improve the contiguity of the *P. fragariae* genome through the assembly of a greater amount of repeat rich sequence. Though this assembly still fell short of the estimated chromosome number of 10 – 12 (Brasier et al., 1999), at 180 contigs, it is a significant improvement over the previous assemblies that utilised solely short read technologies and produced relatively fragmented assemblies, comprised of >1,000 contigs each (Gao et al., 2015; Tabima et al., 2017). The increase in size of the assembly presented (91 Mb), suggests that the assembly includes an increased amount of repetitive sequence, indicating that it represents a more complete genome assembly than previous attempts. Although the assembly presented here was larger than previously reported assemblies, it is similar in size to the related Clade 7b species *P. sojae* (95 Mb; Tyler et al., 2006). This study produced assemblies of an additional ten isolates of *P. fragariae* and three isolates of the closely related raspberry pathogen *P. rubi* with short read technology. These assemblies were of similar sizes and contiguity to those previously published (Gao et al., 2015; Tabima et al., 2017).

All assemblies produced from PacBio and Illumina sequencing were annotated and putative effector genes were predicted. The gene model totals are likely inflated, due to the use of a greedy approach to effector gene prediction, to ensure the capture of all possible virulence genes. ApoplastP appeared to over-predict effectors, likely due to statistical issues arising from the use of a training set far smaller than the query set used in this study (Pritchard and Broadhurst, 2014). Interestingly, twice as many CRNs were predicted in *P. rubi* than *P. fragariae* on average (average of 118 in *P. rubi* and 67 in *P. fragariae*), though this was not consistent for all *P. rubi* isolates. In comparison to *P. sojae*, the BC-16 isolate was predicted to possess approximately 50% more RxLRs and twice as many CRNs from RNA-Seq guided gene models (Tyler et al., 2006). This difference may have been due to improvements in prediction strategies, as the number of CRNs was similar to those predicted for the Clade 1 species *P. cactorum* (Armitage et al., 2018). However for RxLR effectors this difference is likely explained by the greedy approach taken for gene prediction in this study. Following the validation of gene models by RNA-Seq data, the likely overprediction of effector genes was also shown by the low percentage of these classes of genes showing evidence of expression.

This work also allowed the resolution of *P. fragariae* race schemes between different countries, Canadian race 1 is equivalent to UK race 1, Canadian race 2 is equivalent to UK race 3 and both Canadian race 3 and USA race 4 is equivalent to UK race 2. This provided further support for the proposed gene-for-gene model of resistance in this pathosystem (van de Weg, 1997a). However, construction of orthology groups for all isolates of *P. fragariae* and *P. rubi* did not show the presence of proteins unique to the races UK1, UK2 and UK3. Additionally, the identification of variant sites, with the BC-16 genome acting as a reference, indicated there were only private variants present in UK2, with none identified in UK1 or UK3. None of these variants were in proximity to genes thought to be involved in pathogenicity. Additionally, analysis of population structure confirmed the previously described species separation of *P. fragariae* and *P. rubi* (Man in ’t Veld, 2007; Tabima et al., 2018) and identified a subpopulation consisting of the isolates of *P. fragariae* of UK1-2-3 on which further investigations focused.

Transcriptome analyses led to the identification of a strong candidate for *PfAvr2*, which was shown to be highly expressed in all *in planta* BC-16 timepoints, compared to evidence of no, or very low levels of expression in any BC-1 (UK1) or NOV-9 (UK3) samples. The strongest candidate for *PfAvr3* was also shown to be differentially expressed between races, but not as highly expressed in NOV-9 as observed for putative *PfAvr2*. RT-qPCR confirmed the *PfAvr2* results and additionally showed expression of *PfAvr2* in the other sequenced UK2 isolate, A4. The assay showed high levels of variability, which was due to variation between biological replicates, likely the result of the inability of the inoculation method to control the quantity of zoospores inoculating an individual plant. Further *in planta* experiments and RT-qPCR of additional isolates are required to investigate the observations of putative *PfAvr3*.

We propose that silencing of putative *PfAvr2* in races UK1 and UK3, enables those isolates to evade recognition in *Rpf2* possessing plants, such as ‘Redgauntlet’, but not in plants possessing ‘*Rpf1*’ or ‘*Rpf3*’, respectively. Silencing of putative *PfAvr3* also enables races UK1 and UK2 to evade recognition in *Rpf3* possessing plants, as long as they also do not possess *Rpf1* and *Rpf3*.

Whilst the exact mechanism of putative *PfAvr2* and *PfAvr3* silencing was not identified in this study, long read sequencing of a race UK3 isolate (NOV-9) identified a single SNP, 14 kb downstream of the stop codon of putative *PfAvr2* and variation in expression of transcription factors was observed in the RNA-Seq data. This indicated that epigenetic modifications could explain the observed transcriptional variation. Transcriptional silencing of effectors is a known mechanism that *Phytophthora* spp. employ to evade the activation of host *R*-gene-mediated immunity. Investigations of the EC-1 clonal lineage of *P. infestans* revealed a variation of the ability of isolates to cause disease on potato plants possessing the *Rpi-vnt1.1* gene. It was shown, in the absence of genetic mutations, that differences in the expression level of *Avrvnt1* were detected that correlated with virulence (Pais et al., 2018). Recent work in *P. infestans* and *P. sojae* has shown evidence of adenine N6-methylation (6mA) alongside a lack of evidence of 5-methylcytosine (5mC) DNA methylation, highlighting that 6mA methylation is an important epigenetic mark for the regulation of gene expression in *Phytophthora* spp. (Chen et al., 2018). It is also possible that chromatin modifications may be involved in controlling the expression differences, as demonstrated in *P. infestans* (van West et al., 2008; Chen et al., 2018). Investigation of these possibilities, while not achievable in the current study is a clear direction for future research.

Nearly all *P. fragariae* isolates investigated in this study were collected from around Canada between 2001-2012. The SCRP245 isolate was identified as an intermediate between the UK1-2-3 population and the population represented by the BC-23 and ONT-3 isolates. SCRP245 could represent a rare hybrid of the two populations. However, due to the small sample size of isolates, the low number of SNP sites available for this analysis and the fact it was isolated in 1945 in the UK, at least 55 years before the other isolates, it appears more likely that SCRP245 represents a separate population but the data available were unable to resolve this fully. Further race typing with differential strawberry genotypes is required to ascertain the relationship between isolates of Canadian races 4 and 5.

Races of asexual species have been shown to evolve by the stepwise accumulation of mutations (Del Mar Jiménez-Gasco et al., 2004). One such example is the successive evolution of multiple pathotypes in a single clonal lineage of the wheat pathogen *Puccinia striiformis* f. sp. *tritici* in Australia and New Zealand (Steele et al., 2001). In comparison, our data do not indicate that *P. fragariae* has undergone simple stepwise evolution of effectors, but we rather postulate that some lineages of *P. fragariae* have been present for long periods of time in nature, evident by the large number of SNP differences and well supported branches and that the emergence of races (e.g. UK1-2-3) is fairly recent, possibly as a result of the *R*-genes deployed in commercial strawberries. The increased selection pressure on *P. fragariae* to overcome these genes, or the break-up of ‘wild’ R gene stacks upon hybridisation of octoploid strawberry species, may have led to very rapid evolution of races, in this case through epigenetic silencing of gene expression, to evade the *R*-genes present in common cultivars. Substantial further sampling from multiple geographic regions would be required to fully decipher population structure in the lineages of *P. fragariae* and the resistance status of wild octoploid *Fragaria* species. We predict that this would lead to the observation of other geographically distinct lineages of genetically similar individuals of *P. fragariae* but with similar differences in pathogenicity on strawberry. The implications of these findings highlight the potential adaptability of *P. fragariae* to modify effector expression to evade host resistance and the threat of the emergence of new races. Future strawberry breeding efforts must deploy cultivars with multiple resistance genes to mitigate against the rapid adaptation of *P. fragariae*. Identifying resistance genes that recognise the conserved core RxLRs identified in this study would enable broad-spectrum resistance to this pathogen to be deployed that would be effective against multiple races. One of the conserved highly expressed RxLRs was shown to have homology to *P. sojae Avr1b* and so identifying homologous *R*-genes to *Rsp1b* in strawberry could be a future avenue of work to provide resistance against multiple races.

In conclusion, we have shown for the first time that within a distinct subpopulation of *P. fragariae* isolates, displaying remarkably low levels of polymorphisms, the ability to cause disease on a range of differing strawberry cultivars was associated with variation in transcriptional levels rather than being due to sequence variation, similar to reports in *P. infestans* (Pais et al., 2018). This work therefore has implications for the identification of putative avirulence genes in the absence of associated expression data and highlights the need for detailed molecular 576 characterisation of mechanisms of effector regulation and silencing in oomycete plant pathogens. In addition, this study presents a large amount of data, including an improved, long read assembly of *P. fragariae* alongside a collection of resequenced isolates of *P. fragariae* and *P. rubi*, and transcriptome data from multiple isolates that is a valuable resource for future studies.

## AUTHOR CONTRIBUTIONS

RH, CN and TA devised the study. RH, CN and JD co-supervised TA’s PhD study, within which some of this work was undertaken. TA performed the experimental work, with contributions from CN. AA assisted with the development of genome assembly, annotation and orthology pipelines. MS developed elements of the variant calling pipeline and structure analysis with TA. HB performed Nanopore and MiSeq sequencing. JT, BK, BT and NG provided RNA-Seq reads of *P*. *rubi* for genome annotation. TA drafted the manuscript with input from CN, JD and RH. All authors read and approved the submission.

## ACKNOWLEDGEMENTS

The authors gratefully acknowledge the East Malling Strawberry Breeding Club for access to strawberry material, to Dr. Andrew R. Jamieson for the *P. fragariae* isolates as well as Dr. David E.L. Cooke for the *P. rubi* isolates and *P. fragariae* SCRP245. The authors also gratefully acknowledge Prof. Sophien Kamoun, Dr. Eric van de Weg and Dr. Thijs van Dijk for useful discussions. In addition, we wish to thank the following people for the help received in the preparation and setting up of experiments Dr. Helen M. Cockerton, Dr. Laura A. Lewis and Mr. Joseph Hutchings, as well as the members of RH’s group. Work was carried out under the terms of DEFRA Plant Health Licence 6996/221427 held by RH/CN.

## DATA AVAILABILITY STATEMENT

The datasets, including all assemblies and annotations, generated for this study are available on NCBI GenBank as part of BioProjects PRJNA396163 and PRJNA488213 with accession numbers of SAMN07449679 – SAMN07449692. Raw sequencing reads have been deposited in the NCBI SRA, DNA-Seq reads are available with accession codes SRR7668085 – SRR7668100 and *P. fragariae* RNA-Seq reads are available with accession codes SRR7764607 – SRR7764615. *P. rubi* RNA-Seq reads are available with the accession codes SRR10207404 – SRR10207405.

## SUPPLEMENTARY MATERIAL

### SUPPORTING TABLES

**SUPPORTING CANDIDATE TABLE | Gene names of all genes identified as putative candidate avirulence genes for *Phytophthora fragariae* UK1, UK2 and UK3.** Gene names are listed with respect to the isolate for which they are described as candidate avirulence genes.

**SUPPORTING ANNOTATION TABLE | Details of all predicted genes in *Phytophthora fragariae* UK2 isolate BC-16.**

**SUPPLEMENTARY FIGURE S1.**
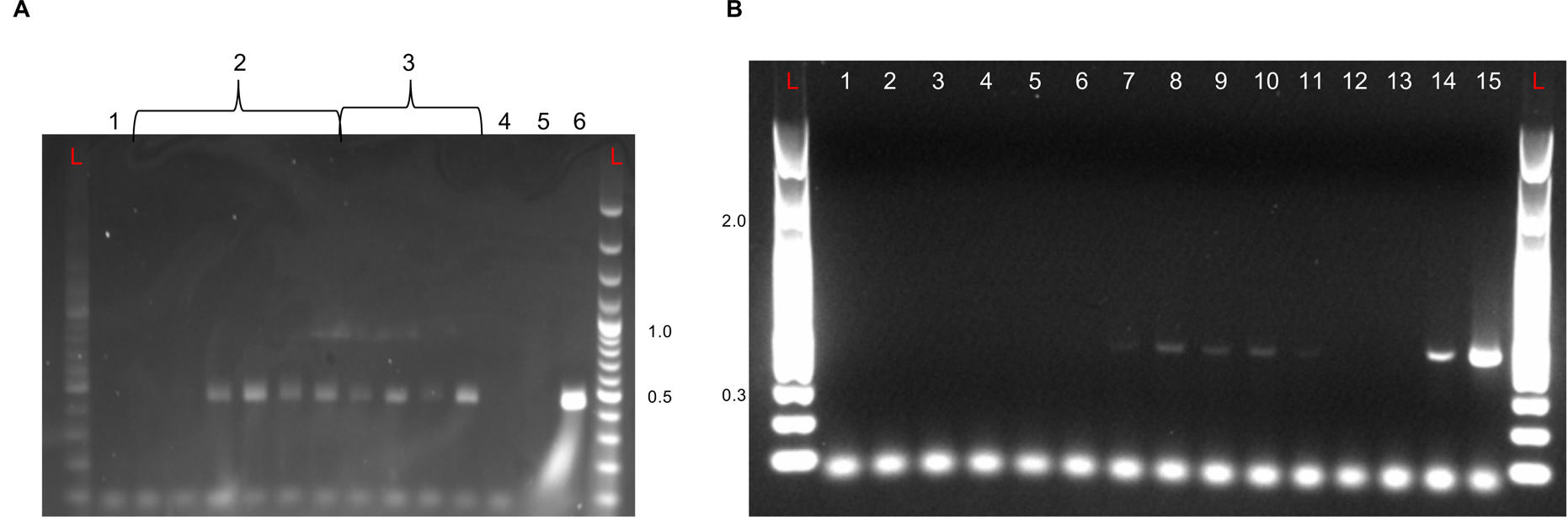
Agarose gel electrophoresis of RT-PCR reactions on representative samples from inoculation time course experiments on the ‘Hapil’ cultivar of *Fragaria* × *ananassa*. **A**: Gel of samples from an inoculation time course experiment with the BC-16 isolate of *Phytophthora fragariae*. **L**: 100 bp Plus Ladder (ThermoFisher Scientific, Waltham, MA, USA) sizes listed in kb. 1: Mock inoculated *Fragaria* × *ananassa* cultivar ‘Hapil’ plant. **2**: Time course of inoculated plants with stream water used as the flooding solution. Time points from left to right: 24 hpi, 48 hpi, 96 hpi, 144 hpi, 192 hpi and 240 hpi. **3**: Time course of inoculated plants with Petri’s solution used as the flooding solution. Time points from left to right: 24 hpi, 48 hpi, 96 hpi and 144 hpi. **4**: dH2O control. **5**: ‘Flamenco’ gDNA control. 6: BC-16 gDNA control. **B**: Gel of samples from an inoculation time course experiment with the BC-1 and NOV-9 isolates of *P. fragariae*. L: 100 bp plus ladder from New England Biolabs, sizes listed in kb. PCR templates: 1: Uninoculated plant cDNA. 2: NOV-9 12 hours post inoculation (hpi) cDNA. **3**: BC-1 12 hpi cDNA. 4: NOV-9 24 hpi cDNA. 5: BC-1 24 hpi cDNA. **6**: NOV-9 48 hpi cDNA. **7**: BC-1 48 hpi cDNA. **8**: NOV-9 72 hpi cDNA. **9**: BC-1 72 hpi cDNA. **10**: NOV-9 96 hpi cDNA. **11**: BC-1 96 hpi cDNA. **12**: dH2O. **13**: gDNA from a *F*. × *ananassa* cultivar ‘Hapil’ plant. **14**: gDNA from BC-16 mycelium. **15**: cDNA from BC-16 mycelium.

**SUPPLEMENTARY FIGURE S2.**
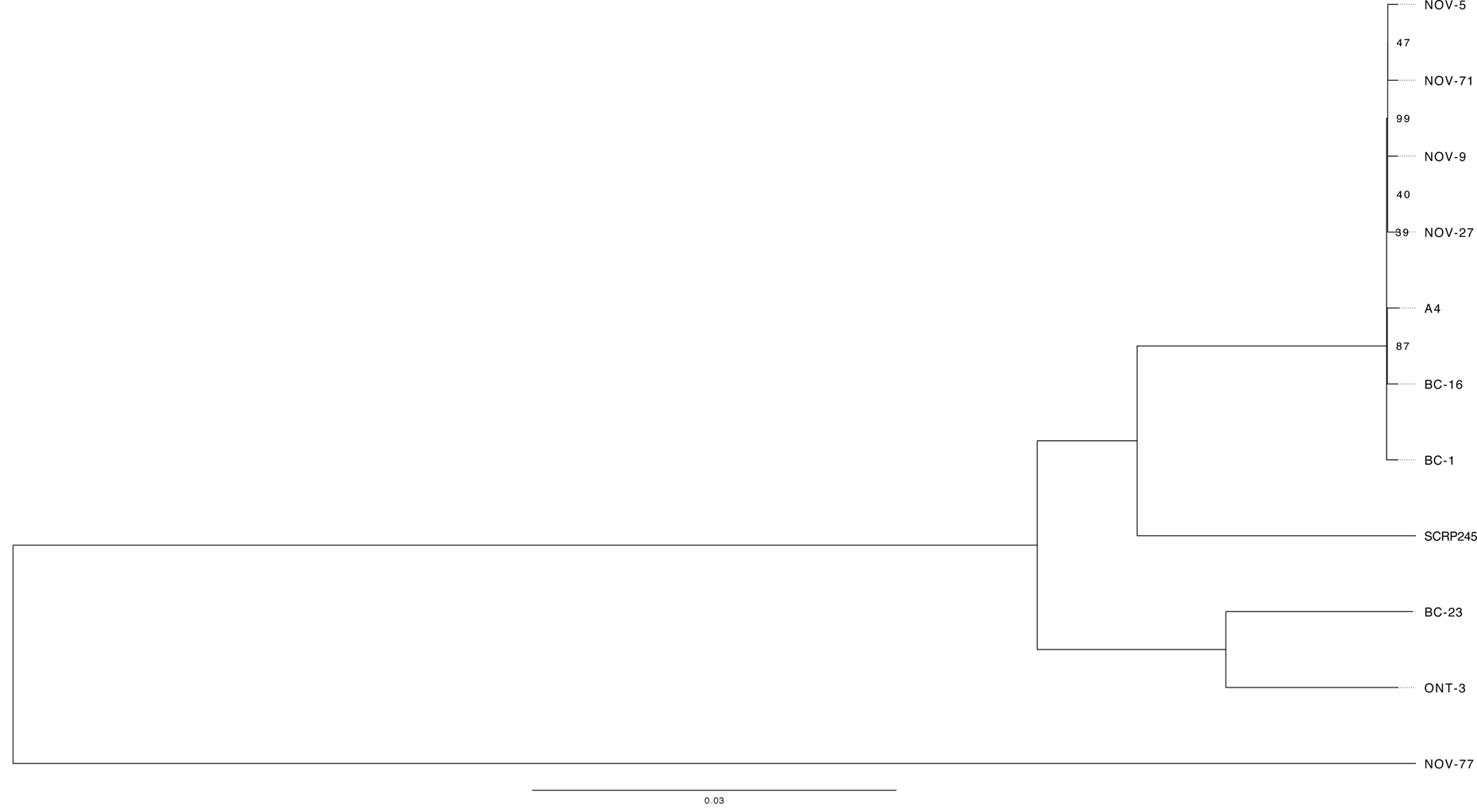
*Phytophthora fragariae* isolates separate into distinct clades. Neighbour joining tree based on high quality, biallelic SNP sites, node labels represent the number of bootstrap replicates supporting the node; only values less than 100 are shown.

**SUPPLEMENTARY FIGURE S3.**
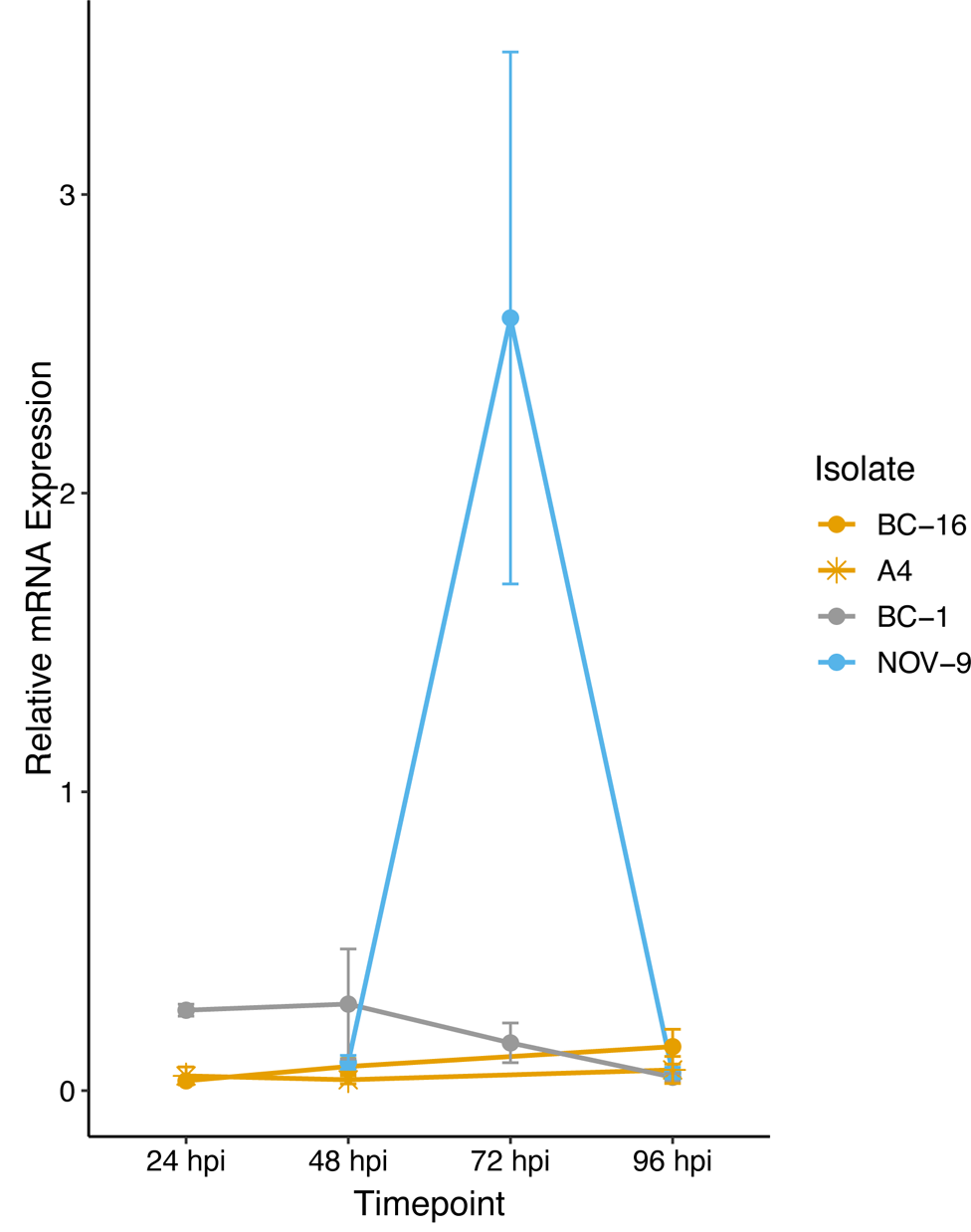
*PfAvr3* candidate PF009_g26276 is differentially expressed in *Phytophthora fragariae* UK-1-2-3 isolates. Quantitative reverse transcription PCR of a candidate for the avirulence gene possessed by NOV-9, but not BC-1, BC-16 and A4 (PF009_g26276.t1). Plots created by the ggplot2 R package (Wickham, 2016) in R version 3.4.3 (R core team, 2017).

**SUPPLEMENTARY TABLE S1.**
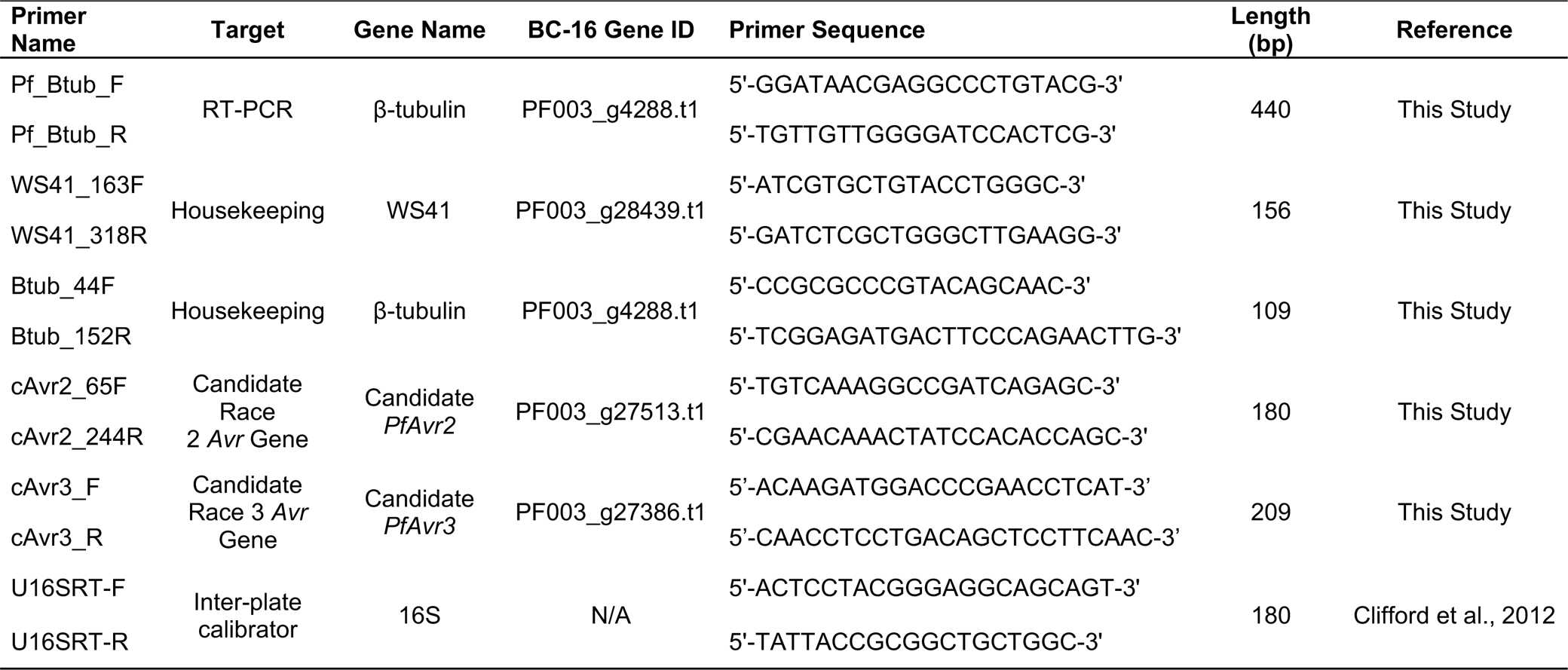
Primers used in this study. Primers supplied by IDT (Leuven, Belgium).

**SUPPLEMENTARY TABLE S2.**
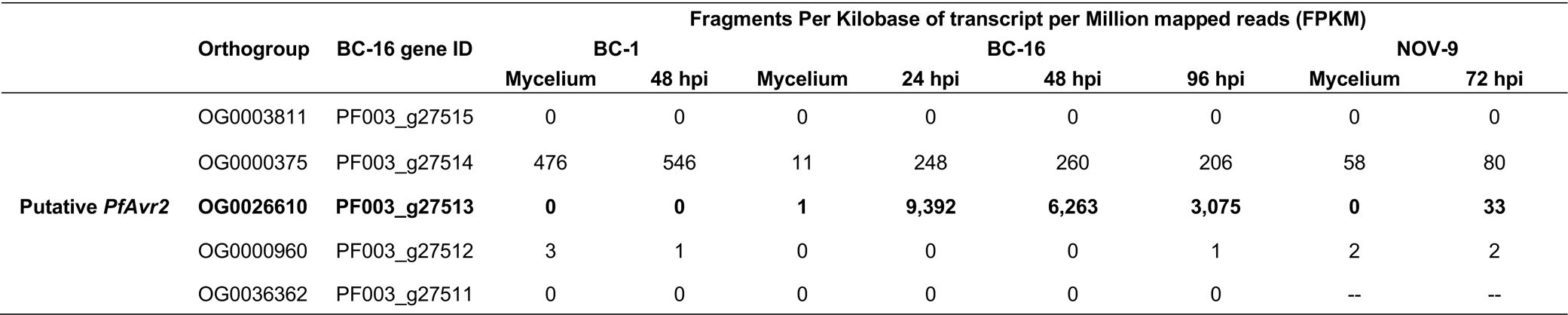
Details of expression of genes surrounding putative PfAvr2 (PF003_g27513).

**SUPPLEMENTARY TABLE S3.**
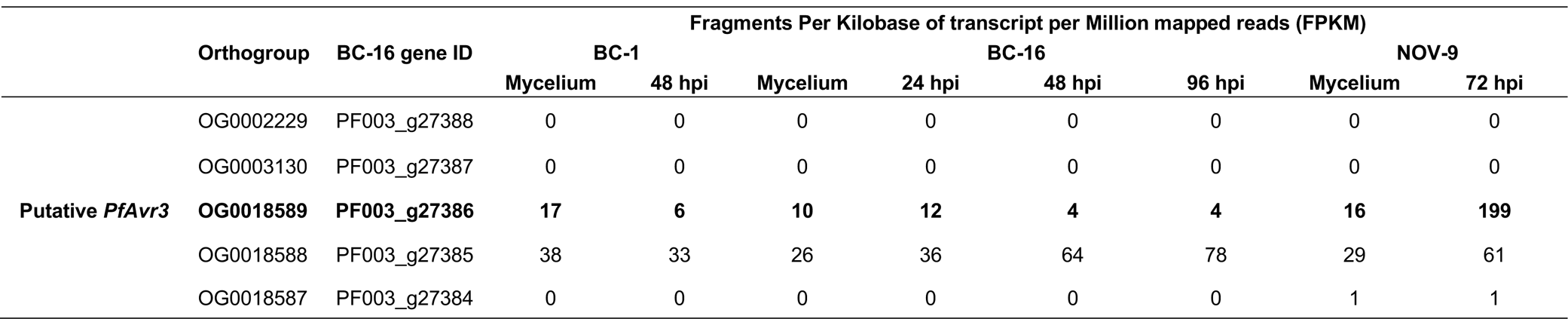
Details of expression of genes surrounding putative PfAvr3 (PF003_g27386/PF009_g26276).

